# Glia-enriched human brain organoids for studying developmental alterations in leukodystrophies

**DOI:** 10.64898/2026.10.01.755898

**Authors:** Elisabeth Mangiameli, Liza M.L. Kok, Clarissa Rosato, Giovanna Cenini, Francesca A. Cupaioli, Roberta Alfieri, Shreenidhi Vitchanthangal Prathivathibayankaram, Polina Oberst, Michael Peitz, Stephanie Dooves, Ivan Merelli, Angela Gritti, Oliver Brüstle, Vivi M. Heine

## Abstract

Leukodystrophies (LKD) comprise mostly inherited white matter disorders whose developmental origins and underlying pathomechanisms remain poorly understood. Progress has been limited by restricted access to human primary tissue and by the inability of animal models to fully recapitulate human neurodevelopment. Human induced pluripotent stem cell-derived neural organoids provide an alternative approach, but most culture systems do not capture later, glia-rich developmental stages relevant to these diseases. Recent studies have shown that appropriate patterning cues and long *in vitro* differentiation can generate neuroglial spheroids comprising multiple neuronal and macroglial populations, including mature oligodendrocytes. Here, we adapt the human oligodendrocyte (hOLS) spheroid platform to establish 3D models of globoid cell leukodystrophy (GLD), 4H leukodystrophy (4H), and Canavan disease (CD). Here we adapt this organoid platform to establish 3D models of globoid cell leukodystrophy (GLD), 4H leukodystrophy (4H), and Canavan disease (CD). Across 150 days of differentiation, hOLS from different genetic backgrounds displayed largely comparable growth kinetics and developmental trajectories, comprising multiple neuroglial cell types with predominantly ventral forebrain and midbrain identities. Comparison of LKD and control hOLS revealed both shared and disease-specific alterations, with prominent transcriptional changes in lysosomal, metabolic, and stress-response pathways. Lineage-resolved pathway analysis further identified signatures across individual LKDs, including mitochondrial and translational stress in GLD glial lineages, disruption of RNA metabolic and cell-cycle pathways in 4H, and prominent astrocyte-centered metabolic dysregulation in CD. Our results establish hOLS as a human 3D *in vitro* platform for studying the impact of LKD-associated genetic variants on early neural development and for dissecting lineage-specific disease vulnerability.

## Introduction

Leukodystrophies (LKDs) are a heterogeneous group of rare genetic disorders affecting the white matter of the central nervous system (Raymond et al., 2017; Wolf et al., 2025). Traditionally classified as hypomyelinating or demyelinating disorders, there are meanwhile more than 50 reported variants of LKDs encompassing a broad spectrum of genetic causes, cellular pathologies, and clinical phenotypes (Ceravolo et al., 2024; van der Knaap & Bugiani, 2017). Clinical presentation varies substantially across and within different LKDs, but these disorders most commonly present in childhood and are characterized by progressive neurological deterioration, including loss of motor function and cognitive decline (Bonkowsky et al., 2010; Nowacki et al., 2022). Despite advances in genetic and pathological characterization, disease-modifying treatments remain unavailable for most leukodystrophies (Nagy et al., 2026; Wolf et al., 2025).

A major challenge in leukodystrophy research is the limited understanding of the cellular and molecular mechanisms underlying disease development and progression (Nowacki et al., 2022). Animal models have provided significant insights into the function of genes associated with different leukodystrophies; however, they often fail to fully recapitulate human pathology (Rutherford & Hamilton, 2019), and species differences hamper the translation of findings from animal models to humans (Semple et al., 2013). Primary human tissue has been essential for characterizing major histological abnormalities associated with different leukodystrophies (van der Knaap & Bugiani, 2017), but the limited sample availability and the predominance of end-stage material hamper investigation of the earliest cellular and molecular events underlying disease development. This is particularly relevant because increasing evidence suggests that several leukodystrophies involve abnormalities that arise before overt myelination and extend beyond oligodendrocytes to affect neural progenitors, astrocytes, and neuronal populations (Nowacki et al., 2022). However, the developmental timing and lineage-specific nature of these alterations remain poorly understood.

Human pluripotent stem cell (hPSC) models provide a powerful approach to investigate leukodystrophies during human neurodevelopment and to identify pathogenic alterations that arise before overt white matter pathology becomes apparent. hPSCs can be derived from patients or genetically engineered to introduce disease-associated gene variants, enabling controlled study of pathogenic mechanisms across neural lineages (Cerneckis et al., 2024). Initial hPSC-based LKD studies mostly relied on 2D mono- and co-cultures (Chanoumidou et al., 2020; Chapleau et al., 2025; Dooves et al., 2023; Lieberman et al., 2022; Mangiameli et al., 2021; Shim et al., 2023; Wang et al., 2013), which cannot fully capture the multicellular interactions underlying white matter pathology. More recently, hPSC-derived 3D organoids comprising multiple neural cell types have been employed to study LKD (Evans et al., 2024; Feng et al., 2023; Ferrer et al., 2026). However, substantial variability between differentiation protocols has complicated cross-study comparisons and the identification of shared disease mechanisms.

Systematic assessment of LKD-associated cellular interactions requires a model system that includes neurons, astrocytes, and oligodendrocytes, which typically emerge at later stages of neural development. Building on the study by Marton et al. (2019), who originally described these cultures as human oligodendrocyte spheroids (hOLS), we generated neuroglial spheroids according to this protocol and retained the acronym hOLS throughout this study. These spheroids comprise multiple co-developing neuronal and macroglial populations, providing a unified platform for modeling leukodystrophies in a developmentally relevant human context. Using time-course analyses combining immunofluorescence, bulk and single-cell RNA-seq, we characterized hOLS derived from two demyelinating LKDs, globoid cell leukodystrophy (GLD) and Canavan disease (CD), and one hypomyelinating LKD, hypogonadotropic hypogonadism and hypodontia (4H) leukodystrophy, alongside healthy and isogenic controls. This design enables direct comparison of convergent and disease-specific pathogenic programs and allows investigation of when and in which neural lineages disease-associated alterations first emerge during human development.

GLD is a demyelinating leukodystrophy caused by pathogenic variants in the *GALC* gene, which encodes the lysosomal enzyme galactosylceramidase, required for degrading galactosylceramide (Suzuki & Suzuki, 1970). Loss of GALC function causes accumulation of the cytotoxic metabolite psychosine (Shin et al., 2016), leading to demyelination, neuroinflammation, and neurodegeneration. Clinically, GLD presents with rapid and progressive neurological decline, causing severe impairment of sensory, motor, and cognitive functions (Feltri et al., 2021).

CD is a demyelinating leukodystrophy caused by mutations in the *ASPA* gene, which impair the breakdown of N-acetyl aspartate (NAA) into aspartate and acetate (Matalon et al., 1988). Resulting NAA accumulation and acetate deficiency have been implicated in disrupted osmotic balance (Baslow, 2002), energy homeostasis (Madhavarao et al., 2005), proper myelin production, and histone acetylation. Histopathological hallmarks include intramyelin vacuolization (Namboodiri et al., 2006) and swollen astrocytes with enlarged mitochondria and abnormally structured cristae (Grønbæk-Thygesen & Hartmann-Petersen, 2024). Clinically, CD presents with progressive visual, cognitive, and psychomotor impairment (Hoshino & Kubota, 2014).

4H is a hypomyelinating leukodystrophy caused by mutations in RNA polymerase III subunits, most commonly the *POLR3A* or *POLR3B* genes (Bernard et al., 2011). Clinically, it presents with developmental delay, dental abnormalities, myopia, progressive motor dysfunction, and mild to moderate intellectual disability (Wolf et al., 2014). Beyond hypomyelination, 4H is characterized by oligodendrocyte loss, abnormalities in neurons and glial cells (Perrier et al., 2021; Vanderver et al., 2013), and widespread axonal damage. *POLR3A* mutations are generally associated with worse outcomes, although severity and progression vary among patients (Wolf et al., 2014).

Although the causative mutations and histopathological features of GLD, 4H, and CD have been described (Bernard et al., 2011; Matalon et al., 1988; Suzuki & Suzuki, 1970; van der Knaap & Bugiani, 2017), the developmental timing and lineage-specific mechanisms underlying disease manifestation remain poorly understood. Here, we establish disease-specific hOLS within a unified experimental framework to investigate the developmental timing and cellular origins of leukodystrophy-associated pathology across human neural development. Using longitudinal single-cell and bulk transcriptomics together with molecular and ultrastructural analyses, we assess the ability of this system to identify disease-associated alterations that emerge at early developmental stages and affect progenitor, neuronal, and glial populations before overt myelination. This resource provides a framework for mechanistic studies of leukodystrophies and for identifying lineage-specific therapeutic vulnerabilities.

## Results

### hOLS undergo progressive maturation and generate major CNS cell types

To establish a standardized, human-based platform for studying LKD pathogenesis, we generated human induced pluripotent stem cell (hiPSC)-derived oligodendrocyte-enriched spheroids (hOLS) based on a published protocol (Marton et al., 2019). This 3D system comprises neuronal, astrocytic, and oligodendroglial lineages that co-develop both spatially and temporally, enabling the assessment of cell-cell interactions at different stages of maturation in a human developmental context. Using hiPSCs from healthy donors (HDs) and LKD patients, as well as gene-edited hiPSC lines, we generated spheroids that were analyzed at different days of differentiation (D) (D50, D100, and D150) by immunofluorescence, bulk RNA sequencing (bulk RNA-seq), single-cell RNA sequencing (scRNA-seq), and electron microscopy (**Fig. 1A**).

**Figure 1.**
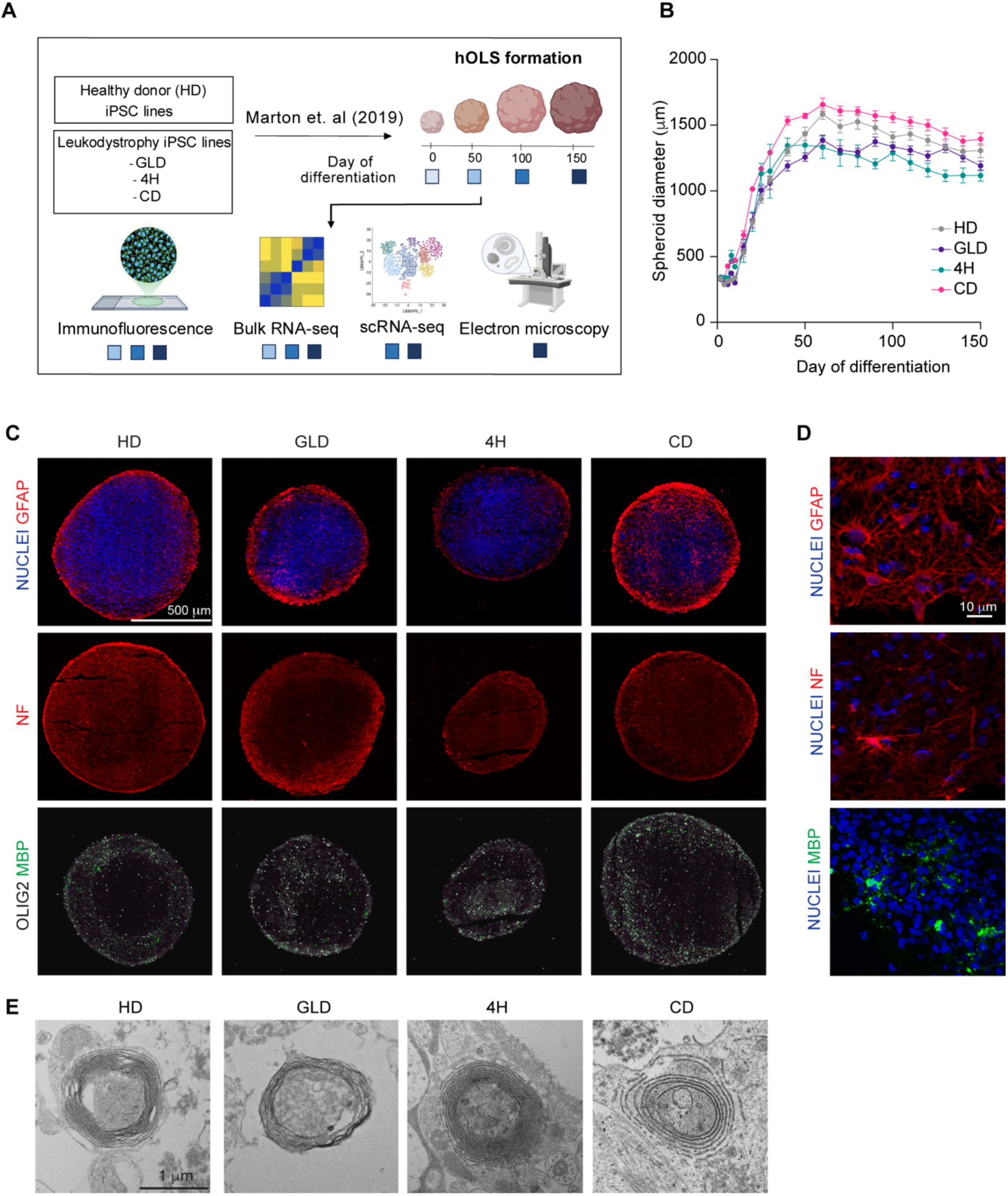
Generation and maturation of oligodendrocyte-enriched human neural spheroids (hOLS) **(A)** Schematic overview of the experimental design. GLD: Globoid Cell Leukodystrophy; 4H: 4H Syndrome or Pol III-related Leukodystrophy; CD: Canavan Disease; RNA-seq: RNA sequencing; scRNA-seq: single cell RNA sequencing. **(B)** Time-course measurements of hOLS diameter. Data are expressed as the mean ± SEM (n= 4-45 hOLS/group/timepoint). **(C)** Representative immunofluorescence analyses on sectioned hOLS at D150 depicting cells positive for GFAP, neurofilament (NF) as well as OLIG2 and MBP. Nuclei are counterstained with Hoechst. Scalebar: 500 μm. **(D)** Magnified images taken from (C). Scalebar: 10 um. **(E)** Representative electron micrographs of D150 hOLS showing myelin sheet formation. Scalebar: 1 μm.

The hiPSC lines used to model GLD, 4H, and CD, together with their respective controls, were generated at different institutes (**Table 1**). The GLD cohort included one patient-derived hiPSC line (GLD 1P) and two CRISPR/Cas9-generated isogenic lines: a *GALC* knock-out (KO) line (GLD 2i) carrying a frameshift mutation and a *GALC* knock-in (KI) line (GLD 3i) carrying the same G553R mutation as GLD 1P. The identity of the gene-edited lines was confirmed by Sanger sequencing (**Fig. S1A, S1B**). All GLD hiPSC lines lacked GALC enzymatic activity (**Fig. S1C**) and accumulated psychosine (**Fig. S1D**), confirming key biochemical hallmarks of GLD. For 4H, we used previously characterized patient-derived hiPSC lines carrying pathogenic *POLR3A* variants, 4H 1P and 4H 2P; (Dooves et al., 2023; Kok et al., 2025). For CD, five ASPA-deficient hiPSC lines derived from two genetic backgrounds and carrying frameshift mutations in exon 1 (CD 1i-5i) were included in the study, along with two isogenic controls (HD4 and HD5). Of those CD 1i-3i were used to generate CD hOLS. ASPA targeting was validated by Sanger sequencing (**Fig. S1E**) as well as ASPA western blot and NAA measurements (**Fig. 4J-K**). Five HD lines were used as controls, including the corresponding isogenic controls for the gene-edited GLD and CD lines (**Table 1**). Together, these lines enabled comparison of patient-derived and gene-edited LKD models within a common hOLS framework. For further details on the iPSC lines used see Table 1.

**Table 1.**
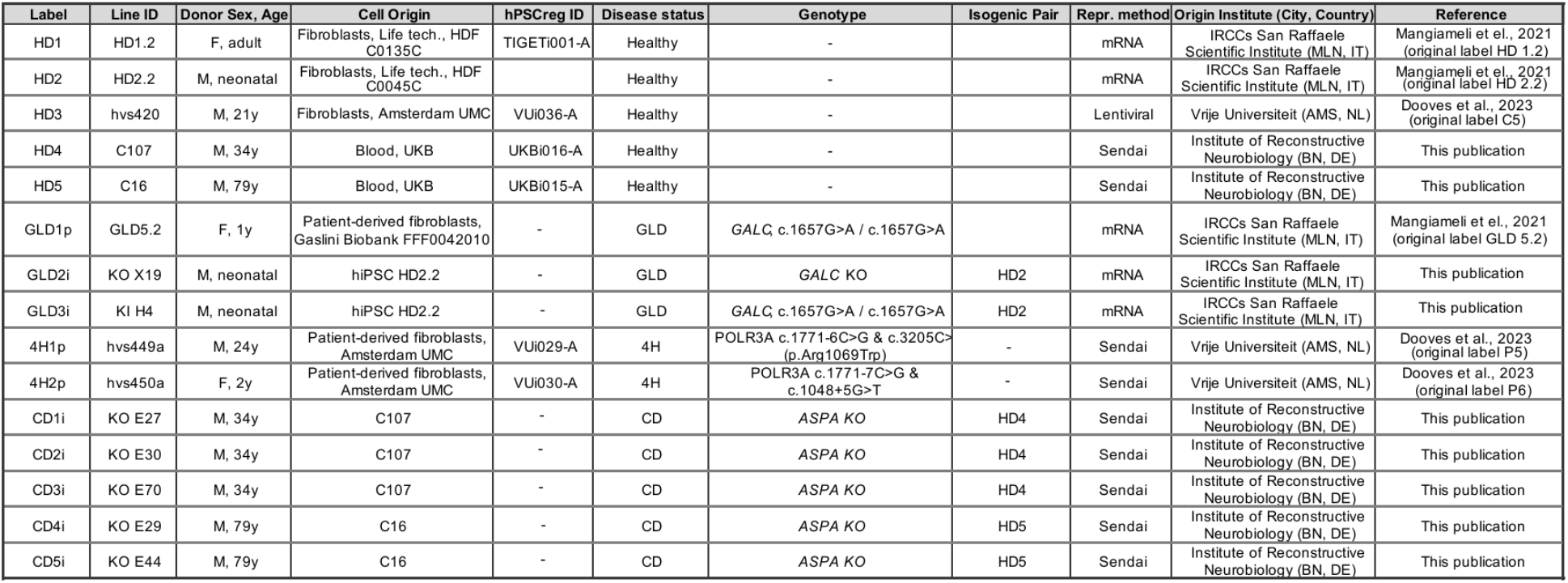
Overview of hiPSC lines used in this study.

To minimize variability and ensure comparability, hiPSC culture and hOLS differentiation were centralized in a single laboratory. The hOLS cultures were regularly monitored for morphological parameters to assess differentiation and overall culture integrity. Diameter measurements revealed an initial phase of rapid growth during early differentiation, coinciding with the period of growth factor supplementation (**Fig 1B**). After 36 days *in vitro*, when growth factor administration ceased, spheroid growth plateaued and remained relatively constant until the end of the culture period (**Fig. 1B**). Despite variability in spheroid diameter across genotypes, LKD hOLS remained within the range observed for HD OLS (**Fig. 1B, Fig. S1F**). At D150, hOLS contained GFAP+ astrocytes, Neurofilament (NF)+ neurons, OLIG2+ oligodendrocyte progenitor cells (OPCs), and MBP+ oligodendrocytes in both HD and LKD conditions, as assessed by immunofluorescence analysis (**Fig. 1C**). MBP+ oligodendrocytes showed a highly ramified morphology, indicative of advanced maturation (**Fig. 1D**). These oligodendrocytes contacted axons and formed myelin sheath-like structures, although not fully compacted, in both HD and LKD samples, as confirmed by electron microscopy (**Fig. 1E**). No necrotic areas were observed until D150 *in vitro*. Together, these findings demonstrate that hOLS generate all three cardinal CNS cell lineages and support oligodendrocyte maturation and myelin formation across healthy and leukodystrophy genotypes, providing a reproducible platform for subsequent molecular and cellular analyses.

### Single-cell transcriptomics reveals diverse neuroglial subtypes and heterogeneous regional identity

To assess the cellular composition and developmental trajectories of hOLS, we performed scRNA-seq on HD and LKD samples at D100 and D150. In HD spheroids, we identified 27 distinct clusters using reference-based annotation against first-trimester developing human brain (Braun et al., 2023), PanglaoDB, and a manually curated signature (see Methods). This analysis revealed a diverse repertoire of cell types spanning progenitor and differentiated neuronal and glial states (**Fig. 2A, 2B, S2**).

**Figure 2.**
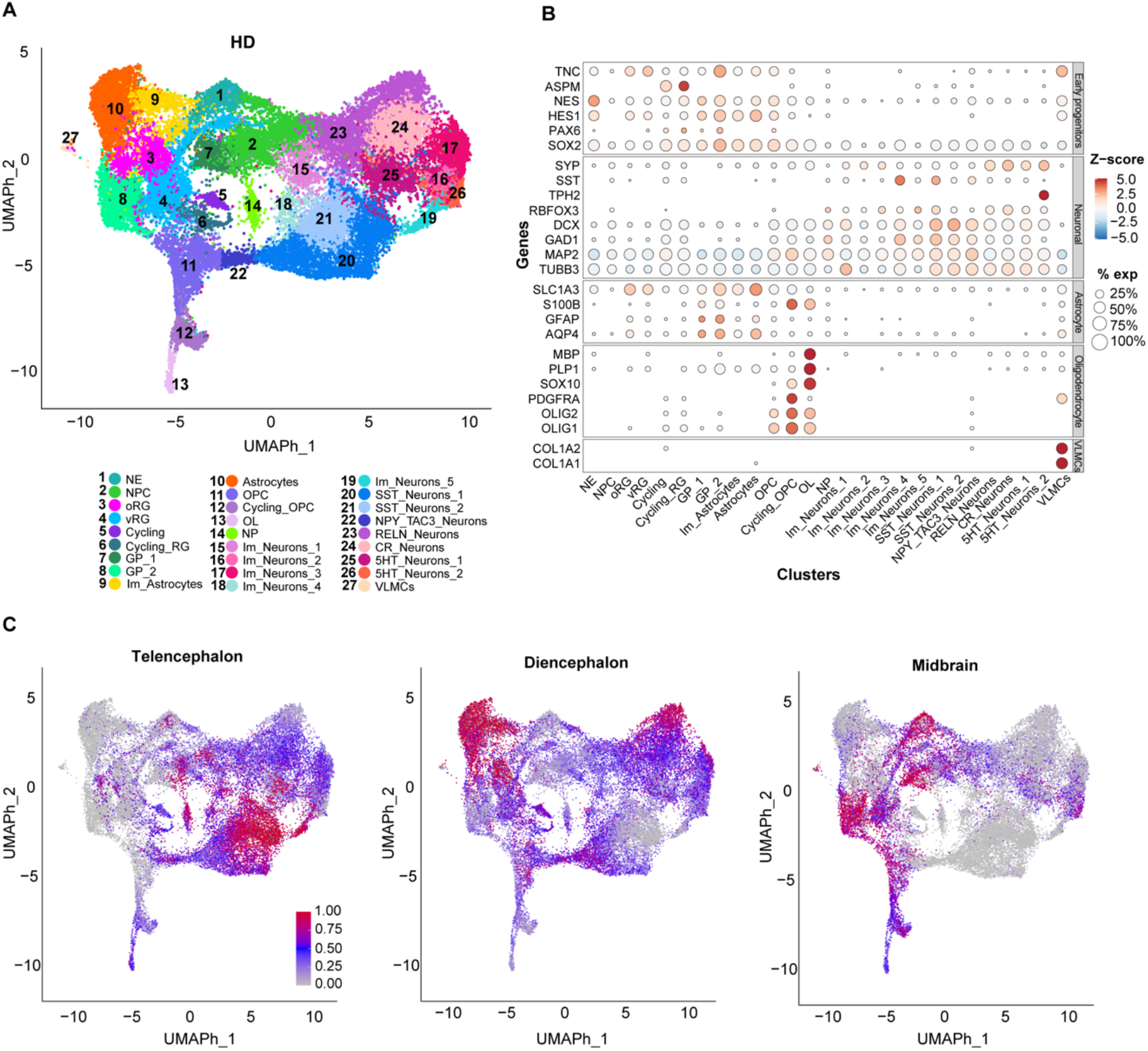
Single cell transcriptomics highlights broad representation of neural cell types in hOLS. **(A)** UMAP plot of scRNA seq data generated with Seurat showing healthy donors hOLS cells at D100 and D150 clustered. **(B)** Bubble plot showing the expression of selected marker genes across single-cell clusters. Bubble size is proportional to the percentage of cells within each cluster expressing a given gene, while color indicates the Z-scored expression level of each gene (blue, low; red, high). **(C**) UMAP plot showing comparison of scRNA-seq analysis of D150 hOLS to Braun fetal human brain atlas dataset. Predicted identity score is depicted according to the color scale shown (grey, low; red, high), 8-14 pcw (post-conceptional weeks) of Braun fetal human brain atlas dataset. From left to right: comparison to telencephalon, diencephalon, midbrain. NE: Neuroepithelial cells; NPC: Neural Progenitor Cells; oRG: Outer Radial Glia; vRG: Ventricular Radial Glia; Cycling: progenitor cells in highly proliferative stage; Cycling_RG: Cycling Radial Glia; GP_1: Glial progenitors; GP_2: Glial progenitors; Im_Astrocytes: Immature Astrocytes; OPC: Oligodendrocytes Progenitors Cells; OL: Oligodendrocytes; NP: Neuronal progenitors; Im_Neurons: Immature neurons; SST_Neurons: Somatostatin neurons; NPY_TAC3_Neurons: Neuropeptide Y/Tachykinin Precursor 3 neurons; RELN_Neurons: Reelin neurons; CR_Neurons: Calretinin neurons; 5HT_Neurons: Serotonergic neurons.; VLMCs: Vascular leptomeningeal cells.

UMAP visualization (**Fig. 2A; HD**) revealed a spatial organization consistent with distinct developmental stages, while specific gene expression signatures (**Fig 2B, S2, S3**) highlighted lineage-specific identities. Neuroepithelial cells (*NES*+/*VIM*+) clustered near radial glia and a distinct proliferative cluster (“Cycling”) that expressed cell-cycle genes but lacked sufficiently specific lineage markers for further annotation. Glial progenitors (*FABP7*+) and astrocytes (*GFAP*+/*AQP4*+) were enriched on the left, with the Glial progenitors 2 (cluster 8) exhibiting ependymal-like features. Oligodendrocyte lineage cells i.e., OPCs (*OLIG2*+/*PDGFRA*+ cluster 11-12) and oligodendrocytes (*MBP*+/*MOG*+, cluster 13) were positioned at the bottom. Neuronal clusters on the right displayed subpallial inhibitory signatures (*DLX2*+/*GAD2*+), including medial ganglionic eminence (MGE)-derived SST+ interneurons expressing *NPY* and *TAC3*, and lateral/caudal ganglionic eminence (LGE/CGE)-like neurons expressing *CALB2* and *RELN* (**Fig. S3A-B**). Additionally, two clusters exhibited serotonergic features, corresponding to precursor (*GATA3*+) and mature (*TPH2*+/*SERT*+) 5-HT neuron populations (**Fig. S3A-B**). Expression of markers for dopaminergic (*TH*), parvalbumin (*PVALB*), cholinergic (*CHAT*), and glutamatergic (*VGLUT2* or *SLC17A6*) neurons was low and scattered, with no detectable expression of *VIP* or *VGLUT1* (data not shown). Interestingly, we also detected a small population of *DCN*+/*LUM*+ cells labelled as vascular leptomeningeal cells (VLMC, **Fig. 2A-B**). No expression of canonical endoderm (*CER1*, *SALL4*), mesoderm (*APLRN*, *PECAM1*), or surface ectoderm (*MSX1*, *KRT8*) markers was detected (**Fig. S4A-D**). Furthermore, pluripotency markers (*POU5F1*, *NANOG)* were undetectable (**Fig. S4E**), supporting hOLS predominant commit toward neural lineages.

To assess how well hOLS recapitulate *in vivo* human brain development, we mapped transcriptomic profiles of HD hOLS onto a developing human brain reference dataset (Braun et al., 2023) using a transfer label approach. Each cell was assigned a prediction score (0–1) reflecting transcriptomic similarity to the corresponding fetal brain population at 5-7 or 8-14 post-conception week (pcw). This analysis revealed broad representation of regional identity with cells allocating to telencephalic, diencephalic and midbrain identities **(Fig. 2C; Fig. S5A**). Whereas neurons from D150 hOLS reflected primarily tel- and diencephalic identities at both mapping stages, astrocytes and oligodendrocytes mapped mainly to midbrain in the pcw 5-7 mapping and to diencephalon in the mapping to 8-14 pcw, indicating temporal differences in regional emergence of these cell types **(Fig. 2C; Fig. S5A**). Importantly, we detected no mapping to fetal cortical (**Fig. S5B**) or spinal cord identities (not shown).

### hOLS undergo lineage-specific maturation along developmental trajectories

We next examined cell maturation dynamics by comparing scRNA-seq of HD hOLS at D100 and D150 (**Fig. 3A**). All cell populations detected at D150 were already present at D100, but their relative abundance changed over time and varied substantially across donors (**Fig. 3B, S6A**). In particular, we observed a clear extension of the UMAP clusters harboring mature astrocytes and oligodendrocytes (**Fig. 3A**). A quantitative depiction of the various cell types across time confirmed an increase in proportion of mature astrocytes and oligodendrocytes between D100 and D150, accompanied by a corresponding decrease in their immature counterparts, while the proportion of immature and mature neurons remained largely stable (**Fig. 3B**). These changes suggest continued maturation of glial populations during extended hOLS differentiation.

**Figure 3.**
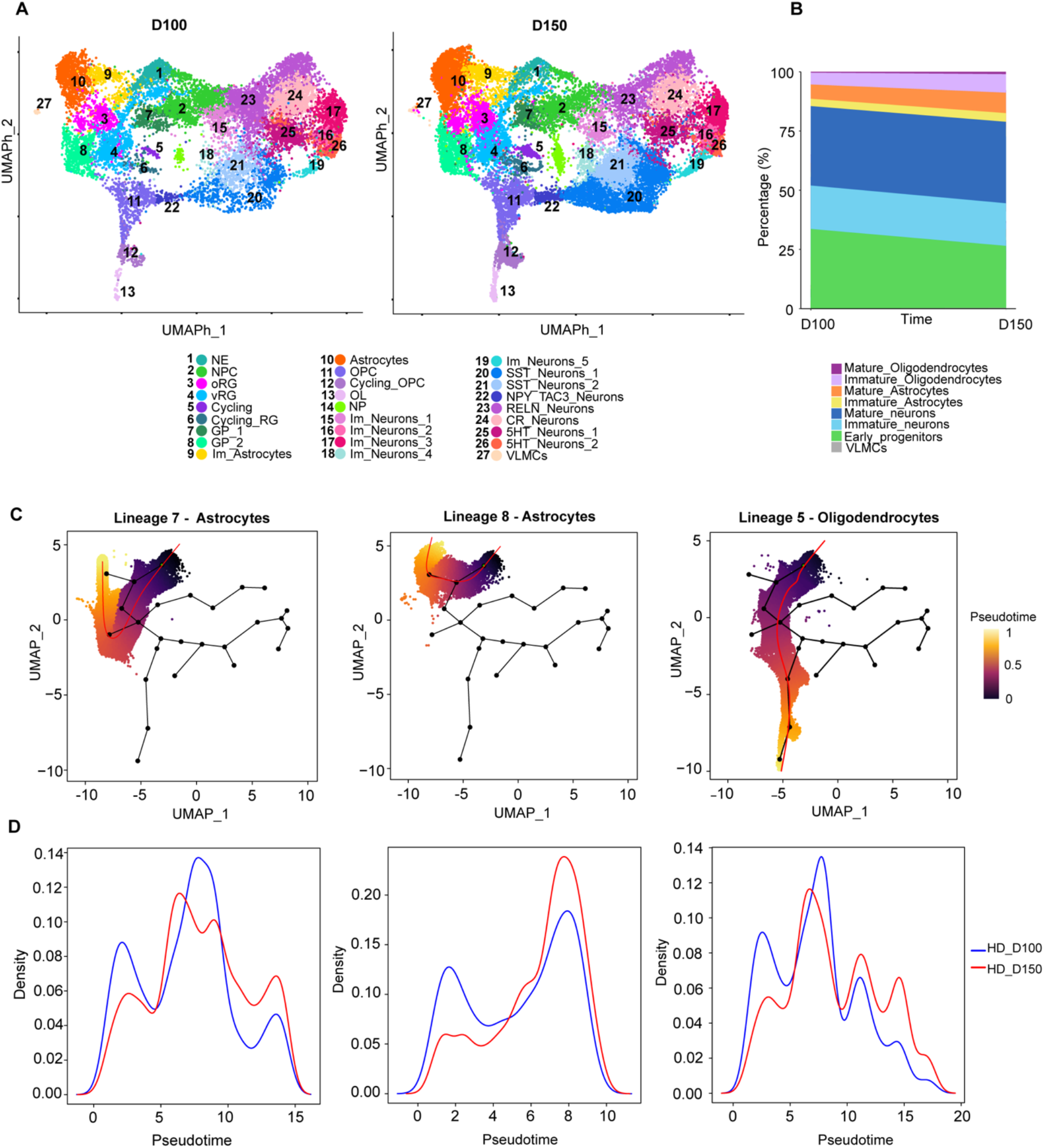
Dynamics of cell differentiation in hOLS. **(A)** UMAP plots showing the clusters identified by scRNA-seq analysis of HD hOLS at D100 and D150. For cluster abbreviations, see the legend to Fig. 2. **(B)** Stacked plot comparing cell-type proportions at the two time points. **(C)** Pseudotime trajectories in HD hOLS. Trajectories were inferred using all genotypes and both time points together. Shown are the two astrocyte trajectories (Lineages 7 and 8) and the oligodendrocyte trajectory (Lineage 5). The color scale indicates pseudotime progression, from early progenitor states (violet) to more mature cell states (yellow). **(D)** Density plots showing pseudotime distributions for the two astrocyte trajectories (Lineages 7 and 8) and the oligodendrocyte trajectory (Lineage 5) in HD hOLS at D100 (blue) and D150 (red).

Considering that hOLS comprise progenitor and differentiated populations with distinct regional identities and developmental states at each time point, inferred pseudotime trajectories may reflect both developmental progression and regional heterogeneity in lineage specification and maturation. With this limitation in mind, we used pseudotime analysis to broadly characterize lineage progression in hOLS. This revealed distinct trajectories originating from the NE cluster and progressing toward terminally differentiated states, which corresponded to oligodendrocytes (Lineage 5), astrocytes (Lineages 7 and 8), and neuronal lineages (Lineages 1, 2, 3, 4, and 6) (**Fig. 3C; S6C**).

Comparison of pseudotime distributions between D100 and D150 revealed lineage-specific maturation patterns (**Fig. 3D**). Cells assigned to the oligodendrocyte (Lineage 5) and astrocyte lineages (Lineages 7 and 8) shifted toward later pseudotime at D150, with four of five donors showing concordant changes for each trajectory (**Fig. 3C-D; S6C**). Neuronal trajectories showed more heterogeneous patterns. Lineage 6 also shifted toward later pseudotime, whereas Lineages 1–3 showed a relative shift toward earlier pseudotime at D150, and Lineage 4 showed little change between time points (**Fig. S6C**). Given the small number of donors, none of these comparisons reached significance at the donor level, and they are therefore reported as descriptive trends. Overall, the pseudotime analysis is consistent with continued progression of glial lineages during extended culture, whereas neuronal trajectories showed less uniform temporal changes.

To validate these changes at the transcriptional level, we analyzed bulk RNA-seq data from HD hOLS. Expression of neuronal markers, including *DCX, GAD1, MAP2, NEUROD1,* and *TUBB3,* and genes related to synaptic functions, namely *SYN1, SYP, and SYT1* remained relatively stable between D50 and D150 (**Fig. S7A-B**). In contrast, glial markers, including *GFAP, MBP, OLIG1, OLIG2, S100B,* and *SOX10,* showed progressive increases in expression (**Fig. S7C)**. Together with the single-cell analyses, these findings support continued gliogenesis and glial maturation during extended hOLS differentiation following an earlier phase of neurogenesis.

In summary, these results demonstrate that hOLS undergo progressive, lineage-dependent maturation, with particularly sustained progression of astrocyte and oligodendrocyte lineages during extended culture. This temporal organization is consistent with the general developmental sequence of neurogenesis preceding gliogenesis and supports the use of hOLS for investigating disease-associated alterations across different stages of neural development.

### hOLS capture disease-relevant features of leukodystrophies

To determine whether hOLS derived from LKD hiPSC lines capture core disease-associated molecular and cellular hallmarks, we integrated single-cell transcriptomics, bulk RNA sequencing, enzymatic activity assays, and metabolic profiling. Principal component analysis (PCA) of bulk RNA-seq data across genotypes and time points (D50, D100 and D150) revealed a clear separation along PC1 (61.1% of variance), which largely reflected differentiation time (**Fig. 4A**). PC2 (7.5% of variance) captured additional inter-sample variability that did not clearly align with disease status, indicating that developmental stage was the major source of transcriptional variation across hOLS.

To determine whether hOLS recapitulate the core molecular and biochemical features of GLD, we integrated bulk and single-cell transcriptomics with lipid and protein analyses (**Fig. 4B–C**). In HD hOLS, bulk RNA-seq revealed a time-dependent increase in *GALC* and *ASAH1* expression (the latter mediating psychosine production via GalCer deacylation) consistent with ongoing glial maturation (**Fig. 4D**). The *GALC* knockout line carrying a frameshift mutation (GLD 3i) showed reduced *GALC* transcript at all time points, likely due to nonsense-mediated RNA decay (Tuladhar et al., 2019), whereas lines carrying the G553R missense mutation showed near-control-level expression. scRNA-seq revealed broad *GALC* and *ASAH1* expression across neuroepithelial and glial cell clusters, with additional expression in selected neuronal subpopulations and VLMCs, with highest *ASAH1* expression in Glial progenitors 1 (**Fig. 4E**). The overall cellular distribution of these transcripts was similar across HD and GLD hOLS. Western blot analysis confirmed the presence of cleaved, enzymatically active GALC protein in HD hOLS at D150, but not in GLD hOLS (**Fig. 4C**). Consistent with loss of GALC activity, mass spectrometry revealed marked psychosine accumulation of in all GLD lines (**Fig. 4B**), while Galactocerebroside (GalCer) levels - primarily produced in oligodendrocytes as indicated by expression of the GalCer-synthesizing enzyme UGT8 (**Fig. S2A**) – remained within the physiological range (**Fig. 4B**). Glucosylceramide (GlcCer) and glucosylsphingosine levels were reduced in GLD hOLS (**Fig. 4B**), suggesting broader disruption of sphingolipid metabolism. Together, these findings show GLD hOLS recapitulate key molecular and biochemical hallmarks of GALC deficiency, including psychosine accumulation (Kobayashi et al., 1988).

For 4H, bulk RNA-seq analysis showed a modest decline in *POLR3A* expression over time in HD hOLSs, whereas the two patient lines displayed different expression patterns. *POLR3A* expression was consistently lower in 4H 1P across all time points, whereas 4H 2P remained closer to HD levels (**Fig. 4F**). The differential reduction in *POLR3A* transcripts between the two patient lines may reflect mutation-specific effects on transcript stability or RNA processing, as pathogenic POLR3A variants have been reported to differentially affect splicing, transcript abundance, and protein function (Kok et al., 2025). In contrast, *POLR3B* expression remained largely stable across samples and differentiation stages (**Fig. 4F**). ScRNA-seq analysis revealed that *POLR3A* and *POLR3B* were ubiquitously expressed across neural lineages, with high expression levels in neural progenitors and selected neuronal subtypes, including SST neurons (**Fig. 4G**). The overall cellular distribution of these transcripts was comparable between HD and 4H hOLS, consistent with a global requirement for RNA polymerase III activity across neural cell types. Together, these findings suggest that POLR3 defects do not reflect restricted lineage expression, raising the possibility that specific neural populations are differentially vulnerable to impaired RNA polymerase III activity.

As for CD, the CD-relevant transcripts *ASPA* and *NAT8L* showed a clear increase over time in bulk RNA-seq data of HD hOLS, (**Fig. 4H**), reflecting maturation of N-acetylaspartate (NAA) metabolism. ScRNA-seq showed that *ASPA* and *NAT8L* transcripts were most abundant in *excluded from **(E)**. For cell-type abbreviations, see the legend to Fig. 2. **(J)** Representative western blot showing absence of ASPA protein in two ASPA-KO hOLS clones at D150; βIII-tubulin (TUBB3) was used as loading control. **(K)** N-acetylaspartate (NAA) levels in culture supernatants from HD and CD hOLS at D150. Each dot represents an individual donor*.

astrocytes, with additional expression detected in glial progenitors and radial glia, whereas NAT8L was also expressed in a subset of immature neurons (**Fig. 4I**). In ASPA KO hOLS, western blot analysis at D150 confirmed the absence of ASPA protein expression in contrast to its robust expression in their isogenic control counterparts (**Fig. 4J**). Consistent with ASPA deficiency, NAA levels at D150 were significantly higher in the culture supernatants of CD hOLS (**Fig. 4K**), reflecting impaired NAA breakdown and confirming a core metabolic feature of CD. These findings show that the hOLS model recapitulates tissue accumulation of NAA and further support a prominent role for astroglial metabolic dysfunction in CD pathogenesis.

**Figure 4.**
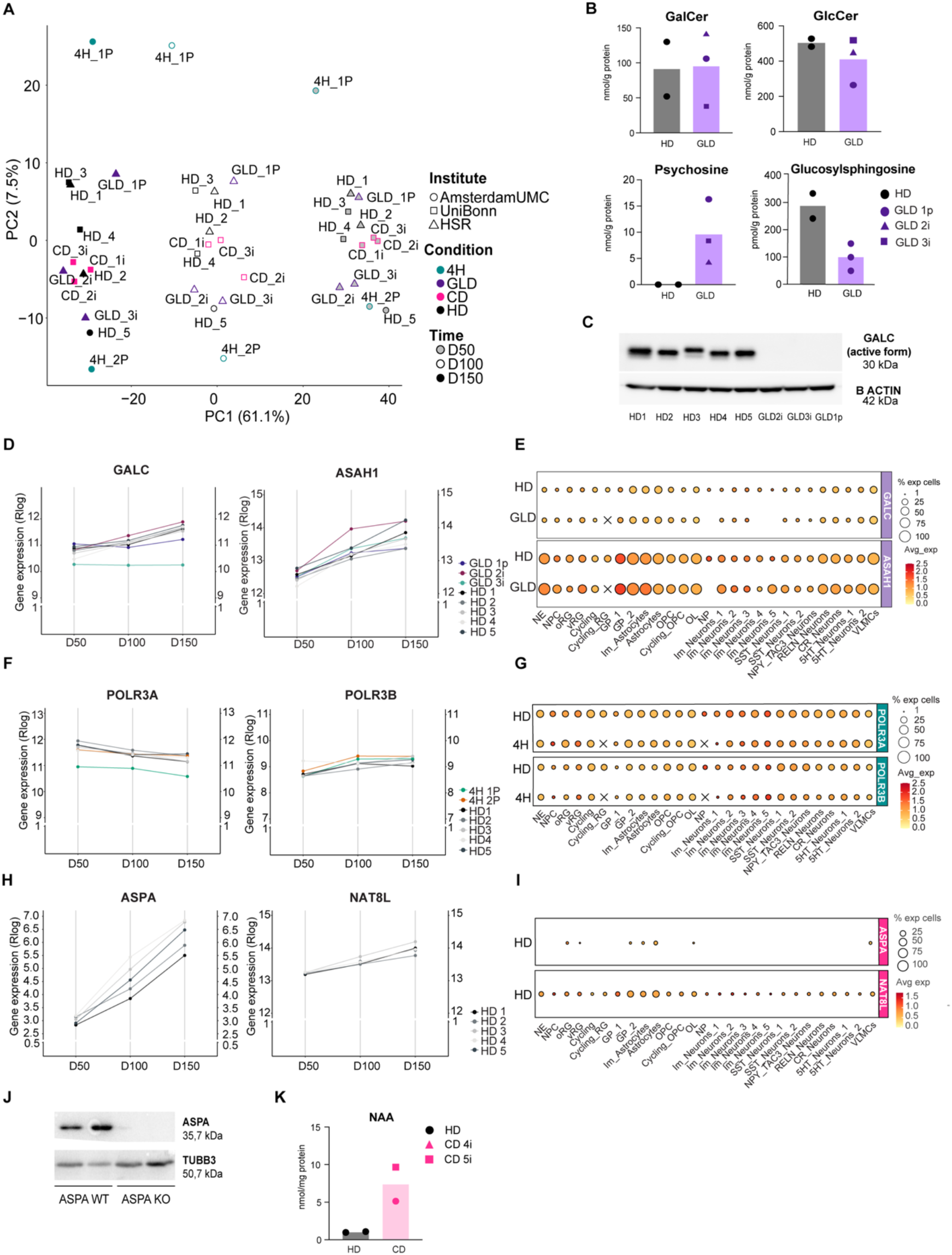
hOLS recapitulate molecular and metabolic hallmarks of leukodystrophies. **(A)** Principal component analysis (PCA) of bulk RNA-seq profiles from hOLS of different disease and control backgrounds at D50, D100, and D150. **(B)** Quantification of galactocerebroside (GalCer), glucosylceramide (GlcCer), psychosine, and glucosylsphingosine by mass spectrometry in HD and GLD hOLS at D150. Data are shown as individual donors. **(C)** Representative western blot of cleaved active GALC protein in HD and GLD hOLS at D150; β-actin was used as loading control. **(D, F, H)** Time-course bulk RNA-seq expression profiles of disease-related genes in HD and LKD hOLS: GALC and ASAH1 for GLD **(D)**, POLR3A and POLR3B for 4H **(F)**, and ASPA and NAT8L for CD **(H)**. **(E, G, I)** Bubble plots showing expression of the corresponding disease-related genes across cell clusters identified by scRNA-seq in HD and GLD **(E)**, HD and 4H **(G)**, and HD **(I)** hOLS. Bubble size represents the percentage of cells expressing the indicated gene and color represents average expression level (yellow, low; red, high). X indicates absence of the respective cluster. GLD 2i was

Together, these results show that hOLS capture disease-relevant molecular and metabolic hallmarks across genetically distinct leukodystrophies while providing access to their expression across different stages of neural development.

### Shared cellular alterations across leukodystrophy hOLS

To investigate cellular alterations across genetically distinct leukodystrophies, we compared HD and LKD hOLS at D100 and D150 using immunofluorescence and scRNA-seq (**Fig. 5**). Immunofluorescence quantifications showed that OLIG2- and MBP-positive areas varied across hOLS but was comparable between HD and LKD conditions (**Fig. 5A**). GFAP abundance progressively increased over time in both HD and LKD hOLS and appeared more pronounced in LKD conditions at D150 (**Fig. 5A**), whereas GABA immunoreactivity was also moderately increased in LKD hOLS at D150 compared with HD conditions (**Fig. 5A**). These findings could suggest alterations in glial and neuronal markers across LKD conditions, although quantitative interpretation was limited by the high cell density and heterogeneous cellular representation across hOLS sections.

**Figure 5.**
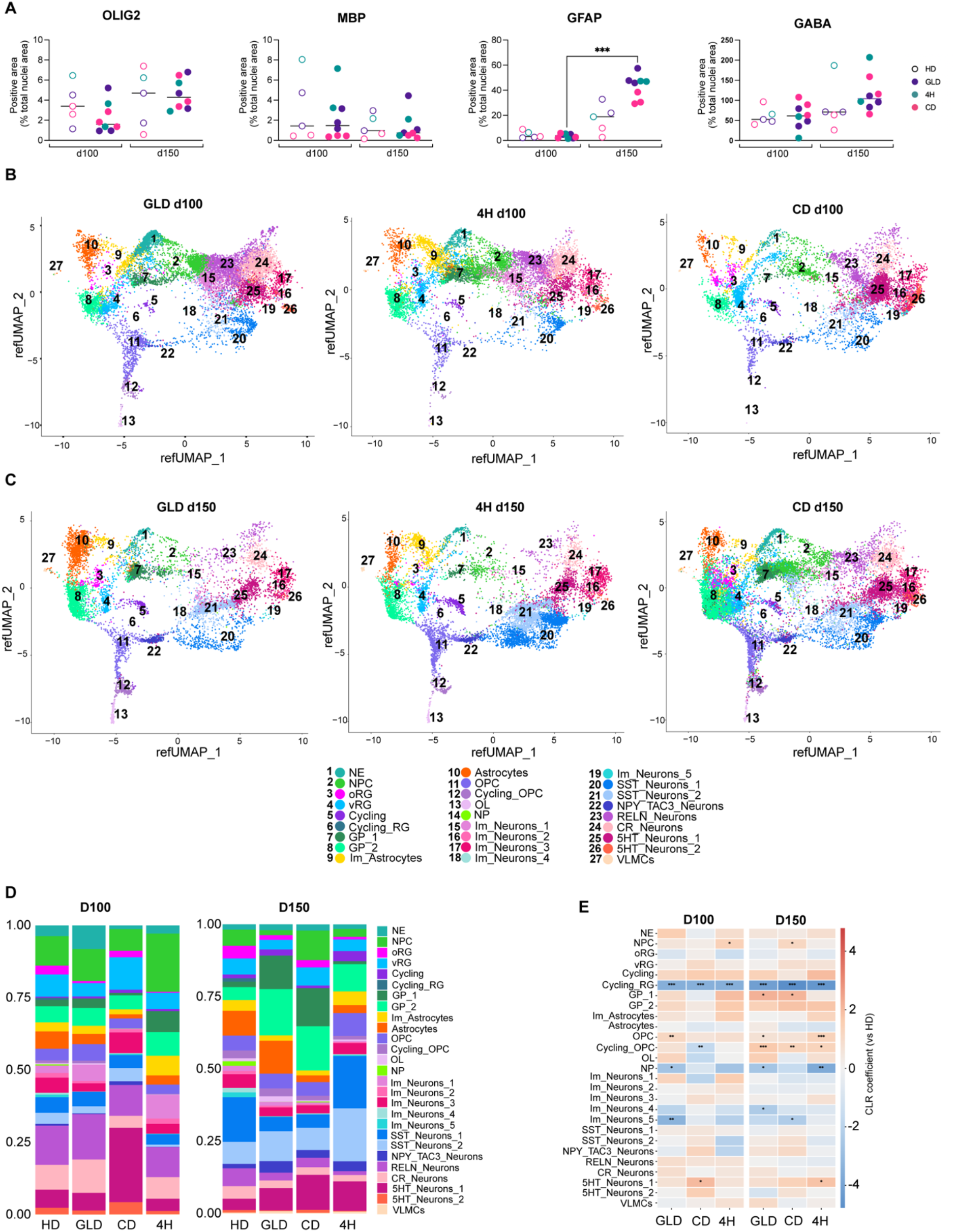
Cell type composition across leukodystrophy hOLS. **(A)** Quantification of immunofluorescence staining in hOLS sections at D100 and D150. OLIG2-, MBP-, GFAP-, and GABA-positive areas were normalized to nuclear area. Each dot represents an individual donor (n = 22–74 sections/group). Statistical analysis was performed using the Kruskal–Wallis test followed by Dunn’s multiple-comparison test; ***p < 0.001. **(B, C)** UMAP representations of cell clusters identified by scRNA-seq analysis of GLD, 4H, and CD hOLS at D100 **(B)** and D150 **(C)**. **(D)** Stacked bar plots showing the relative proportions of cell clusters identified by scRNA-seq in HD and LKD hOLS at D100 and D150. **(E)** Heatmap of linear model coefficients for cell-type composition in GLD, CD, and 4H versus HD hOLS at D100 and D150, estimated from centered log-ratio (CLR)-transformed values. Asterisks indicate statistical significance (*FDR < 0.1; **FDR < 0.05; ***FDR < 0.01). For cell-type abbreviations, see the legend to Fig. 2.

ScRNA-seq analysis provided a broader and more detailed view of cellular composition across LKD hOLS. UMAP visualization showed that the major neural cell populations identified in HD hOLS were also represented in GLD, 4H, and CD hOLS at D100 and D150 (**Fig. 5B, C**). Quantification of cluster proportions, however, revealed differences in the relative representation of specific cell populations across disease conditions and developmental stages (**Fig. 5D**). These differences were particularly evident among progenitor, immature neuronal populations and OPCs. Quantitative analysis of selected clusters showed reduced proportions of cycling radial glia in GLD, 4H, and CD hOLS compared with HD, with particularly low representation in 4H and CD (**Fig. 5E**). Neuronal progenitors were also reduced in GLD, while immature Neurons 4 and 5 showed lower representation across LKD conditions, although the magnitude and statistical support of these changes varied between diseases and time points (**Fig. 5E)**. Within the oligodendrocyte lineage, LKD hOLS exhibited a trend towards increased proportions of cycling OPCs and OPCs, particularly at day 150, without a corresponding increase in mature oligodendrocytes. Together, these observations indicate that LKD-associated differences in cellular composition become already visible in progenitor and immature populations during hOLS development.

We then asked whether these alterations were also reflected in the position of cells along the inferred differentiation trajectories. Pseudotime distributions were compared between HD and LKD hOLS using linear mixed models accounting for variability between cell lines. Among the three diseases, GLD hOLS showed the most consistent trend, with cells positioned at earlier pseudotime than HD in six of eight lineages at D100, reaching nominal significance in four of them. Effects in CD and 4H were weaker and less consistent. However, none of these comparisons remained significant after correction for multiple testing. Thus, while the analysis suggested potential disease-associated shifts in selected trajectories, we found no robust evidence for systematic alterations in pseudotime progression across LKD hOLS.

Together, these analyses identify overlapping alterations in progenitor and immature neural populations across leukodystrophy hOLS, indicating that disease-associated differences in cellular composition emerge during early stages of neural development.

### Pathway analysis identifies shared and lineage-specific alterations

To identify molecular pathways associated with leukodystrophy-related transcriptional changes, we performed gene set enrichment analysis (GSEA) on both bulk and scRNA-seq datasets. This integrated approach enabled us to compare global transcriptional changes with their cell type-specific signatures.

GSEA of the bulk RNA-seq data identified both shared and disease-specific pathway alterations across GLD, 4H and CD hOLS relative to HD controls (**Fig. S8**). Although most significantly dysregulated pathways were disease-specific, about 80 pathway terms were altered across all three LKDs at both D100 and D150. Shared upregulated programs included neurogenesis, cell development, and cholesterol metabolism, whereas catabolism, autophagy, and lysosomal function were commonly downregulated. These findings reveal that genetically distinct LKD exhibit partially overlapping developmental and metabolic disturbances.

To determine whether disease-associated transcriptional signatures were already present at an earlier developmental stage, we performed unbiased GSEA at D100 (**Fig. S9A–C**). Among the top upregulated and downregulated terms, GLD hOLS were strongly enriched for translation, oxidative phosphorylation, and inflammatory pathways; 4H hOLS exhibited alterations in RNA processing, translation, and stress pathways; and CD hOLS displayed prominent metabolic and mitochondrial changes. Thus, disease-specific transcriptional alterations arise before overt myelination.

Upon closer inspection, bulk GSEA of GLD hOLS at D150 identified broad dysregulation of developmental, metabolic, and stress-response programs (**Fig. 6A**). Upregulated terms included morphogenesis, development, action potential regulation, and extracellular matrix organization, whereas downregulated terms included lysosomal and autophagy-related pathways, inflammatory and antigen-presentation signatures, response to nitrogen, and PI3K-associated signaling. When we examined these terms at single-cell resolution (**Fig. 6A**), we found that the bulk-level signal was not uniformly recapitulated across cell populations. Particularly, pathways related to action potential and shared developmental programs were mainly upregulated in immature neurons, while downregulated in mature neurons. Similarly, lysosomal and autophagy-associated pathways were not globally downregulated at the single-cell level; instead, they were upregulated in multiple clusters, including NPCs, radial glia, and immature neurons (clusters 1, 3, and 4). This divergence is not uncommon in organoid studies as pathway-level trends observed in bulk RNA-seq may result from shifts in cellular composition or averaging across heterogeneous populations, and are not always directly superimposable onto single-cell profiles, also due sensitivity and resolution differences (Atamian et al., 2021; Lim et al., 2022). Inflammation-related terms also varied by cell type, being downregulated in oRG and serotonergic neurons (cluster 2), but upregulated in astrocytes, OPCs, OLs, somatostatin neurons (cluster 1), and RELN and CR neurons. Thus, bulk-level enrichment patterns were not uniformly reproduced across individual cell populations, likely reflecting the contribution of both cellular composition and opposing lineage-specific transcriptional responses.

**Figure 6.**
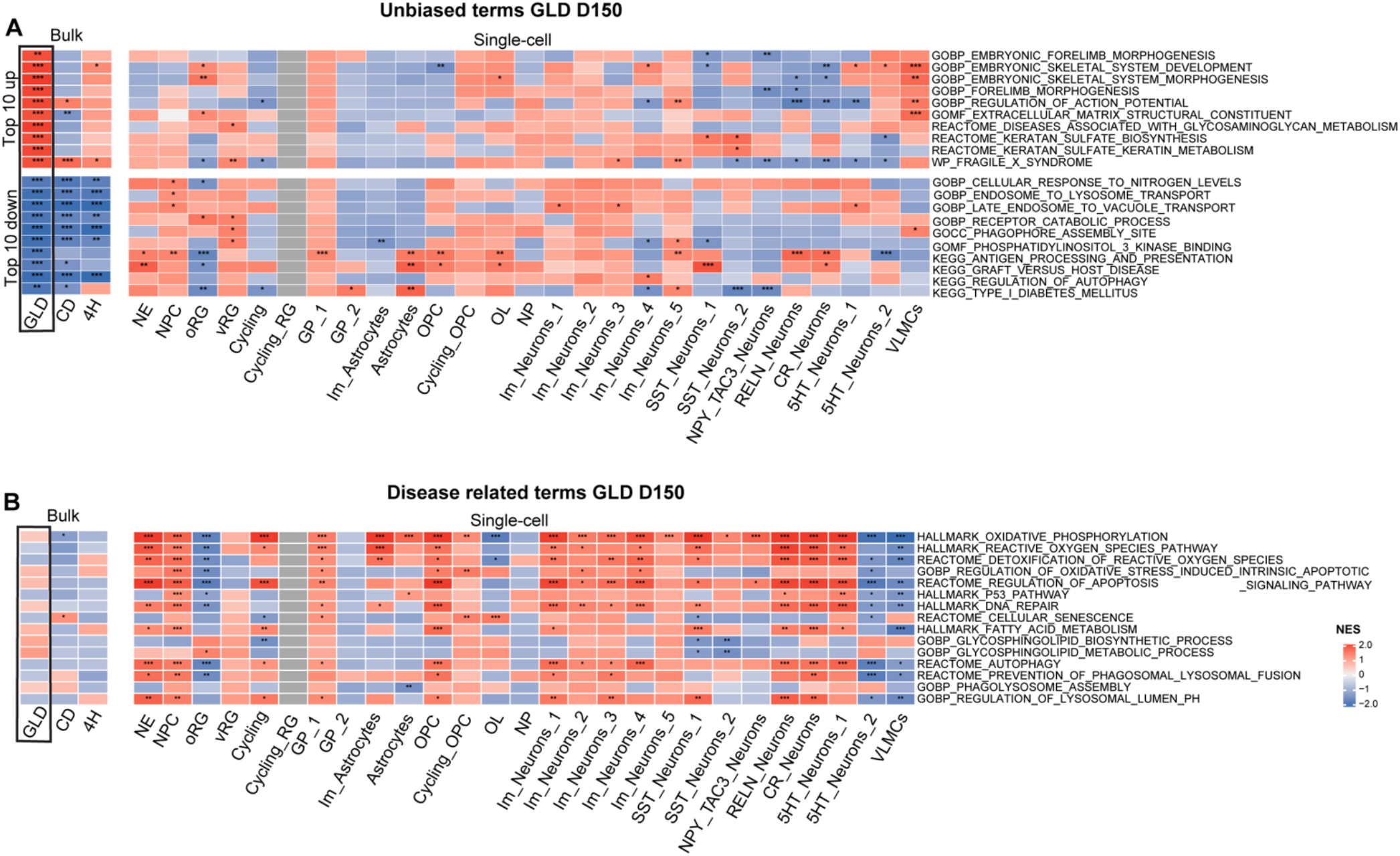
Lineage-specific pathway alterations in GLD hOLS. **(A)** Heatmap showing normalized enrichment scores (NES) of the top 10 upregulated and downregulated GSEA terms identified by bulk RNA-seq analysis of GLD versus HD hOLS at D150 (boxed column). Corresponding NES values for the same pathways are shown for 4H versus HD and CD versus HD bulk RNA-seq comparisons, as well as across cell populations identified by scRNA-seq analysis of GLD versus HD hOLS at D150. **(B)** Heatmap showing NES of selected disease-relevant GSEA terms in bulk RNA-seq comparisons of GLD, 4H, and CD versus HD hOLS (boxed columns indicate GLD bulk RNA-seq) and across cell populations identified by scRNA-seq analysis of GLD versus HD hOLS at D150. The color scale represents NES values, with red indicating positive enrichment and blue indicating negative enrichment. Statistical significance is indicated as follows: *p < 0.05, **p < 0.01, ***p < 0.001. For cell-type abbreviations, see the legend to Fig. 2.

Analysis of GLD-relevant pathways showed modest changes in glycosphingolipid metabolism but marked upregulation of fatty acid metabolism, particularly in OPCs (**Fig. 6B**). Lysosomal and autophagy pathways were significantly upregulated in most clusters at both time points, potentially reflecting a compensatory response to GALC loss and lysosomal dysfunctions. Oxidative phosphorylation was broadly upregulated at D100 (**Fig. S9D)**, whereas at day 150 mature oligodendrocytes and some mature neurons showed reduced oxidative phosphorylation and ROS-detoxification signatures (**Fig. 6B**). Apoptosis- and senescence-associated pathways were also elevated across multiple GLD cell types (**Fig. 6B**). Together, these findings indicate cell type-specific stress responses in GLD, involving lysosomal, metabolic, mitochondrial, apoptotic, and senescence-associated pathways.

In 4H hOLS, bulk RNA-seq at D150 revealed strong upregulation of pathways related to forebrain neuron generation, regionalization, cortical differentiation, GABAergic interneuron differentiation, and neuron fate commitment (**Fig. 7A**). In scRNA-seq, these programs showed pronounced lineage-specific differences, with enrichment in oRG cells but downregulated in NE cells, Im_Neuron_1, and SST neuron populations (**Fig. 7A**). Downregulated bulk pathways included intracellular signaling and endosome transport, while single-cell analysis further showed opposing lineage-specific changes in trafficking-related pathways. Targeted pathway analysis in 4H revealed widespread alterations in early progenitors, glial progenitors, immature neurons, and mature neuronal populations, including serotonergic neurons (**Fig. 7B**). Already by D100, translation and electron transport chain pathways were upregulated across many clusters, particularly in interneuron-associated populations, although directionality varied by lineage (**Fig. S9B, E**). Hedgehog-related and hypoxia-response pathways also showed heterogeneous enrichment patterns. Overall, these results indicate that 4H leukodystrophy involves early and persistent dysregulation of RNA metabolic, neurodevelopmental, and stress-response pathways, with particularly strong effects in progenitor and neuronal lineages.

**Figure 7.**
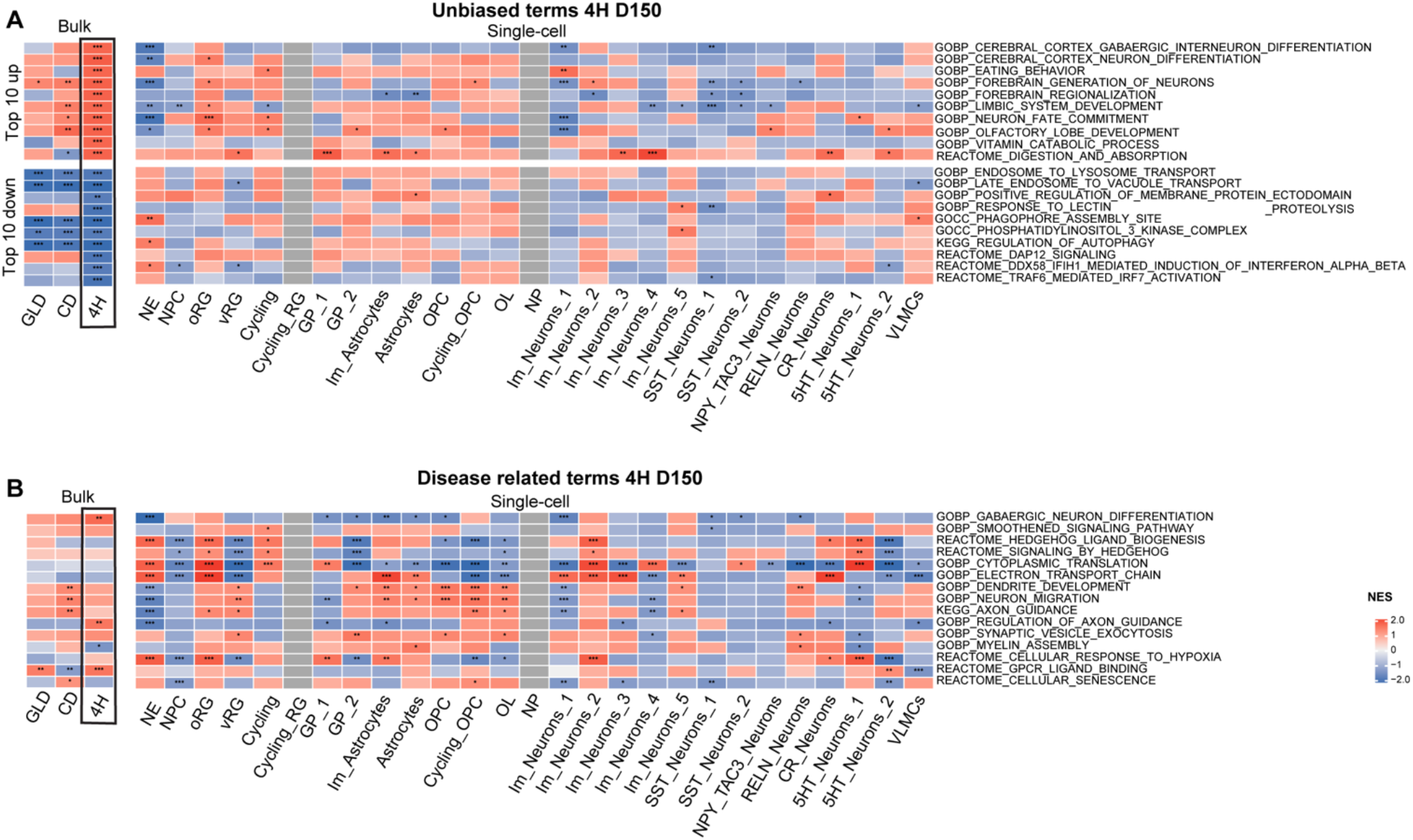
Lineage-specific pathway alterations in 4H hOLS. **(A)** Heatmap showing normalized enrichment scores (NES) of the top 10 upregulated and downregulated GSEA terms identified by bulk RNA-seq analysis of 4H versus HD hOLS at D150 (boxed column). Corresponding NES values for the same pathways are shown for GLD versus HD and CD versus HD bulk RNA-seq comparisons, as well as across cell populations identified by scRNA-seq analysis of 4H versus HD hOLS at D150. **(B)** Heatmap showing NES of selected disease-relevant GSEA terms in bulk RNA-seq comparisons of GLD, 4H, and CD versus HD hOLS (boxed column indicates 4H bulk RNA-seq) and across cell populations identified by scRNA-seq analysis of 4H versus HD hOLS at D150. The color scale represents NES values, with red indicating positive enrichment and blue indicating negative enrichment. Statistical significance is indicated as follows: *p < 0.05, **p < 0.01, ***p < 0.001. For cell-type abbreviations, see the legend to Fig. 2.

In CD, bulk RNA-seq at D150 showed upregulation of pathways associated with mitotic processes, including chromatid segregation and spindle microtubule organization (**Fig. 8A**). ScRNA-seq refined this finding by showing that proliferation signatures were especially enriched in cycling radial glia cells and cycling OPCs (**Fig. 8A**), consistent with altered glial dynamics in CD (Baslow & Guilfoyle, 2009; Mattan et al., 2010). Increased postsynaptic density-related transcripts were also observed in several neuronal subtypes, OPCs, and immature astrocytes, suggesting broader changes in synaptic organization and neuron-glial signaling. Downregulated pathways were more heterogeneous and included responses to nitrogen levels and other metabolic processes, particularly in OPCs **(Fig 8A)**. Analysis of selected CD-relevant pathways revealed pronounced cell type-specific changes in oxidative phosphorylation and mitochondrial metabolism. These pathways were significantly downregulated in immature astrocytes, GP_2, OPCs, and cycling OPCs at both D100 (**Fig. S9F**) and D150 (**Fig. 8B**), whereas these were upregulated in GP_1, radial glia, astrocytes, NPCs **(Fig S9C, Fig. 8B)**. Similar metabolic changes were observed in certain neuronal populations, which could suggest that the effects of ASPA deficiency and resulting NAA accumulation extend beyond glial lineages. We also observed alterations in histone modification and chromatin remodeling pathways. Furthermore, at D100, HAT-mediated acetylation and HDAC-mediated deacetylation were upregulated across several glial populations, consistent with increased chromatin remodeling during early disease stages (Grønbæk-Thygesen & Hartmann-Petersen, 2024; Singhal et al., 2017) **(Fig. S9F)**. By D150, this pattern was largely reversed **(Fig. 8B)**, which could point to temporal changes in epigenetic regulation during hOLS maturation. Together, these findings show that CD hOLS exhibit pronounced glial metabolic dysfunction that may extend to neurons as well as impairment of epigenetic processes relating to acetylation.

**Figure 8.**
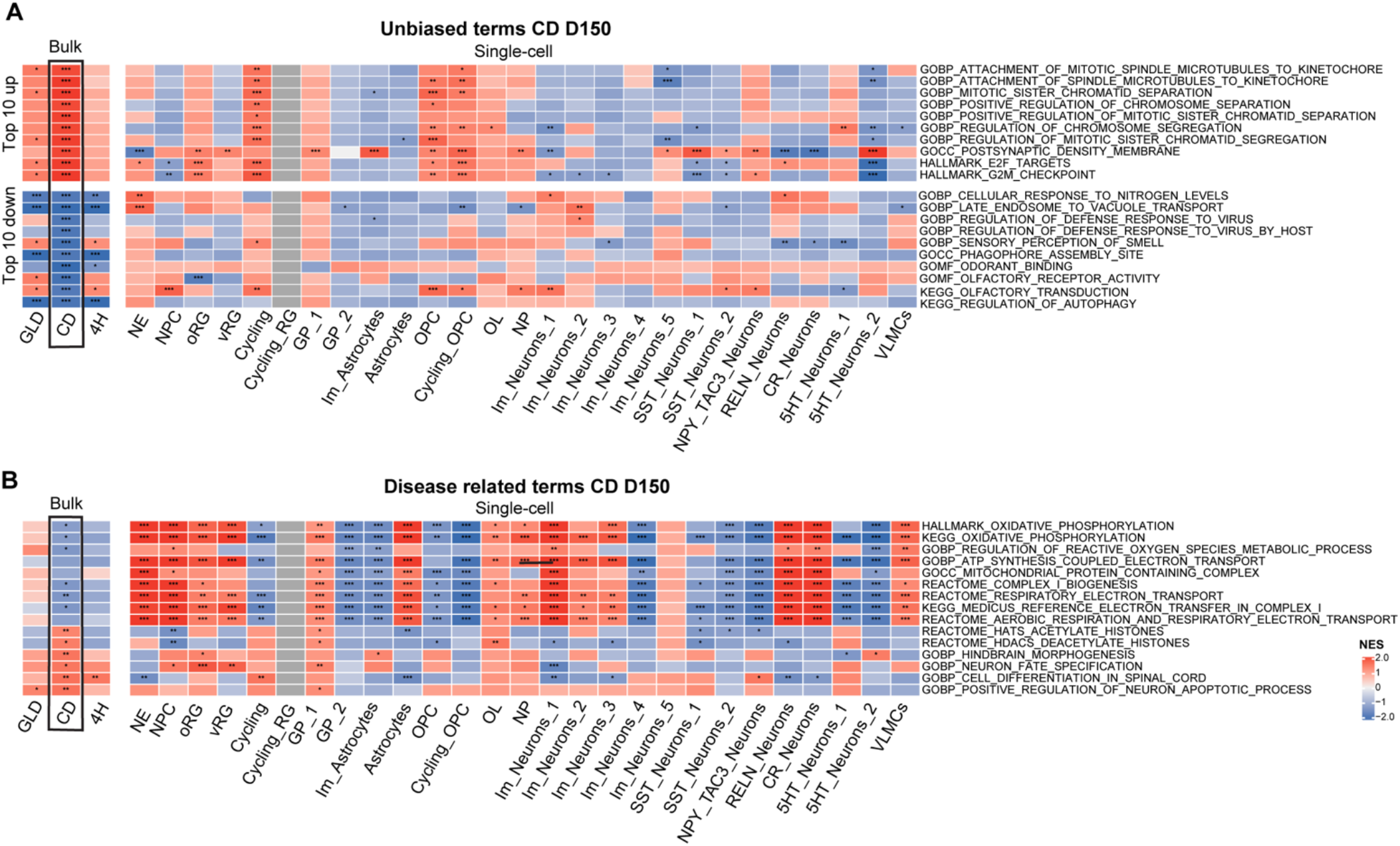
Lineage-specific pathway alterations in CD hOLS. **(A)** Heatmap showing normalized enrichment scores (NES) of the top 10 upregulated and downregulated GSEA terms identified by bulk RNA-seq analysis of CD versus HD hOLS at D150 (boxed column). Corresponding NES values for the same pathways are shown for GLD versus HD and CD versus HD bulk RNA-seq comparisons, as well as across cell populations identified by scRNA-seq analysis of 4H versus HD hOLS at D150. **(B)** Heatmap showing NES of selected disease-relevant GSEA terms in bulk RNA-seq comparisons of GLD, 4H, and CD versus HD hOLS (boxed column indicates 4H bulk RNA-seq) and across cell populations identified by scRNA-seq analysis of 4H versus HD hOLS at D150. The color scale represents NES values, with red indicating positive enrichment and blue indicating negative enrichment. Statistical significance is indicated as follows: *p < 0.05, **p < 0.01, ***p < 0.001. For cell-type abbreviations, see the legend to Fig. 2.

In summary, our analyses show that genetically distinct leukodystrophies exhibit both overlapping pathway alterations and strongly lineage-dependent transcriptional responses, many of which are already detectable at D100.

### Core enrichment genes highlight lineage-specific pathway contributors

To further investigate the gene-level contributors underlying disease-associated pathway enrichment, we examined core enrichment genes from selected disease-relevant pathways at D100 and D150 (**Fig. S10-12**). For each leukodystrophy, pathways were selected based on their relevance to the primary disease mechanism or due to unexpected enrichment across bulk and single-cell analyses. Heatmaps of core enrichment genes highlighted the genes contributing to pathway-level signature across different cell populations and developmental stages.

In GLD, we investigated two pathways of interest: autophagy, a classical pathway often altered in lysosomal storage disorders and skeletal system development, which was unexpectedly strongly upregulated in bulk RNA-seq at D150. Consistent with our GSEA results (**Fig. 6A, B**), the core enrichment genes within REACTOME_AUTOPHAGY showed differential enrichment across cell types (**Fig. S10A**). This signature included genes involved in autophagy initiation (*ATG13, ATG101, ATG16L2*), autophagosome elongation and formation (*MAP1LC3A/C, GABARAPL1, WIPI1/2*), lysosomal maturation and fusion (*LAMP2, LAMTOR3, UVRAG*), and mitophagy/proteostasis (*PINK1, FUNDC1, HDAC6, UBB/UBC*, ubiquitin-conjugating enzymes), indicating heterogeneous regulation of autophagy-related programs across neuronal and glial populations. Core enrichment analysis of GOBP_EMBRYONIC_SKELETAL_SYSTEM_DEVELOPMENT unveiled that the pattern is preferentially associated with glial populations, immature neurons and VLMCs (**Fig. S10B**). The contributing genes included extracellular matrix- and developmental regulators such as *COL1A1* and *WNT5A*, with *COL1A1* upregulation in glial cells and *WNT5A* increased in glial progenitors, astrocytes, and cycling OPCs. Although collagen expression is not a canonical feature of glial cells, it has been reported in astrocytes in injury-related and oxidative stress contexts, where it may contribute to glial scar formation, inflammatory activation, and neuronal dysfunction (Lee et al., 2026). Moreover, increased collagen deposition in the extracellular matrix has been associated with impaired OPC maturation and reduced myelination in other pathological settings (Yamazaki & Ohno, 2025). *WNT5A* is usually linked to a non-canonical WNT state involved in cell migration, polarity, tissue remodeling, and stress adaptation (Kumawat & Gosens, 2016). VLMCs also showed upregulation of several genes within this pathway, although their limited abundance limits interpretation of their contribution to the overall hOLS phenotype.

In 4H, we examined core enrichment genes from the Hedgehog Ligand Biogenesis pathway, which showed significant enrichment and has previously been implicated in 4H (Dooves et al., 2023) (**Fig. S11**). The number of core enrichment genes differed between D100 and D150, reflecting the genes contributing most strongly to the GSEA enrichment signal at each time point rather than differences in overall pathway significance. At D100, 4H hOLS showed a downregulation of *SHH* in glial and progenitor populations, while *GPC5*, which can increase receptor binding of SHH (Li et al., 2011), shows upregulation in astrocytes and subsets of neurons. The strong upregulation of *GPC5* in astrocytes continued at D150 (**Fig S11B**). At D150, several 26S proteasome subunits (*PSMA*, *PSMB*, *PSMC*, and *PSMD* genes) showed an increase in expression in astrocytes and oRG, but were downregulated in GP_2, NPC and vRG. Ubiquitin-related genes, including *UBB*, *UBC*, *UBA52* and *RPS27A*, were downregulated in oligodendrocyte lineage cells and most progenitor populations. Together, these data reveal cell type-specific alterations among genes contributing to Hedgehog-related pathway enrichment in 4H hOLS.

In CD, we examined cell type-specific alterations in KEGG_OXIDATIVE_PHOSPHORYLATION pathway, which revealed distinct temporal and lineage-dependent changes in mitochondrial respiratory chain gene expression at D100 and D150 (**Fig. S12**). At D100, glial progenitors, cycling OPCs, OPCs, but also several neuronal populations exhibited downregulation of genes encoding subunits of complex I (NDUFA, NDUFB, NDUFC, NDUFS, and NDUFV families), complex III (UQCR family), complex IV (COX4I1, COX5A, COX6A1, COX7A2, together with mitochondrial-encoded MT-CO1, MT-CO2, and MT-CO3), and complex V (ATP5 family and MT-ATP6) (**Fig. S12A**). Mitochondrial DNA-encoded genes involved in electron transport, including MT-ND1–MT-ND6 and MT-CYB, were similarly reduced across these oligodendroglial populations. In contrast, astrocytes displayed relatively higher expression of several oxidative phosphorylation genes at D100, a pattern that became more pronounced at D150. At D150, glial progenitors and OPCs continued to exhibit reduced expression of multiple oxidative phosphorylation genes, whereas neuronal populations demonstrated a more heterogeneous expression profile, with exhibiting slightly increased expression of respiratory chain components in selected neuronal subtypes (**Fig S12B**). Notably, the differentially expressed genes comprised both nuclear-encoded and mitochondrial DNA-encoded subunits that collectively constitute complexes I–V of the oxidative phosphorylation machinery, components that are essential for mitochondrial electron transport and ATP generation (Smeitink et al., 2001; Vercellino & Sazanov, 2022). Similar perturbations in mitochondrial metabolism and oxidative homeostasis have previously been reported in ASPA-deficient model systems and Canavan disease, supporting the relevance of mitochondrial dysfunction as a characteristic feature of disease pathology (Francis et al., 2016; Madhavarao et al., 2005).

Together, these analyses refine the pathway-level findings across the three LKDs and identify candidate genes contributing to lineage-specific transcriptional alterations.

### Lineage-resolved pathway analysis reveals disease-specific alterations

GSEA of scRNA-seq datasets revealed distinct cell type-specific molecular signatures across leukodystrophy hOLS **(Fig. 9**).

**Figure 9.**
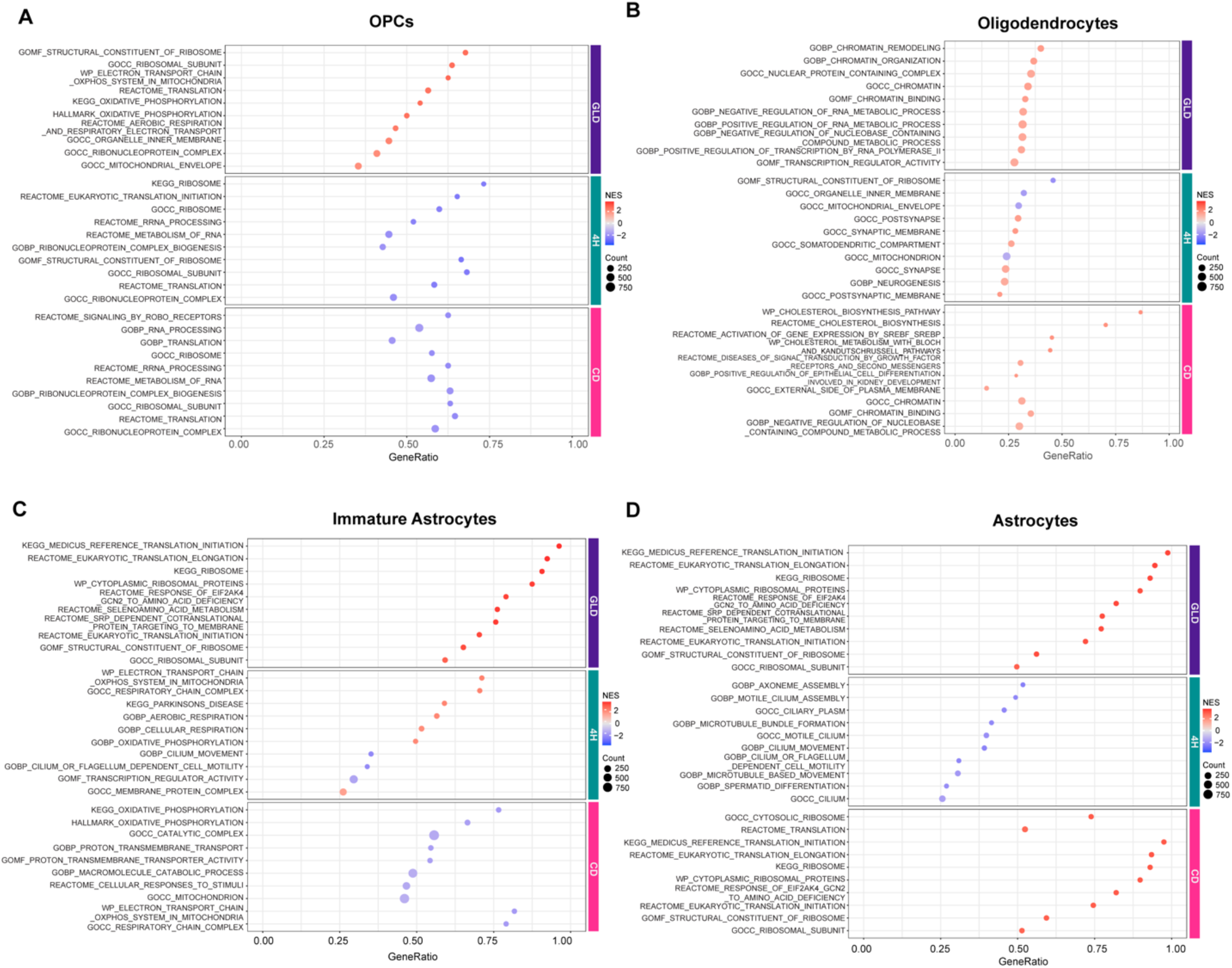
Glial lineage-resolved pathway enrichment signatures at D150. Bubble plots showing the top 10 enriched pathways identified by GSEA of scRNA-seq data from GLD, 4H, and CD hOLS compared with HD hOLS at D150 within: **(A)** Oligodendrocytes Progenitors Cells (OPC), **(B)** Oligodendrocytes, **(C)** Immature Astrocytes, **(D)** Astrocytes. Color indicates the normalized enrichment score (NES), and bubble size represents the number of genes contributing to each pathway.

In GLD hOLS at D150, OPCs showed upregulation of pathways related to chromatin organization and translation (**Fig. 9A**). Similar translation-related signatures were observed in OLs and in immature and mature astrocytes (**Fig. 9B-C**), indicating that alterations in translational programs extend across multiple glial populations. OPCs also showed strong enrichment of mitochondrial pathways (**Fig. 9A**), while genes related to mitochondria and respiration were already perturbed at D100 in OPCs and immature and mature astrocytes (**Fig. S13C, D**). Together, these findings indicate that metabolic and translational alterations are present across multiple glial lineages in GLD and emerge already at earlier stages of hOLS development.

In 4H hOLS, oligodendrocytes showed downregulation of ribosomal and mitochondrial pathways together with upregulation of synaptic development programs at D150 (**Fig. 9B**). OPCs also exhibited reduced translation- and RNA metabolism-related pathways (**Fig. 9A**), suggesting early and persistent defects in RNA-associated programs across the oligodendrocyte lineage. Mature astrocytes displayed downregulation of cilium- and microtubule-associated pathways, whereas immature astrocytes showed upregulated electron transport and respiratory chain pathways (**Fig. 9C, D**). Together with the neuronal defects described in Fig. 7, these findings support broad disruption in RNA metabolism, maturation, and metabolic homeostasis in 4H hOLS.

In CD hOLS, OPCs displayed downregulation of translation and RNA-processing pathways at D150 (**Fig. 9A**), with changes that could point to early disruption of protein synthesis at D100 **(Fig. S13A)**. Oligodendrocytes showed evidence of upregulated cholesterol biosynthesis and metabolism as well as chromatin-associated regulatory pathways at D150 (**Fig. 9B**). At the same time point, immature astrocytes showed reduced oxidative phosphorylation and mitochondrial metabolism, while mature astrocytes were enriched for translation- and ribosome-associated pathways (**Fig. 9C, D**). At D100, both immature astrocyte and astrocyte populations upregulated microtubule organization and cytoskeletal dynamic pathways (**Fig. S13C, D**). Together, these findings reveal temporally and lineage-specific metabolic and transcriptional alterations across oligodendrocyte and astrocyte populations in CD.

Collectively, these analyses reveal distinct lineage-specific transcriptional vulnerabilities across leukodystrophies and identify the neural cell populations contributing to disease-associated pathway alterations.

## Discussion

Leukodystrophies have traditionally been described as disorders of myelin and white matter maintenance; however, the developmental timing and cellular mechanisms underlying disease initiation and progression remain incompletely understood. Here, we employ human hiPSC-derived oligodendrocyte-enriched spheroids (hOLS)(Marton et al., 2019), a spheroid-based brain organoid model, to investigate leukodystrophy pathogenesis across genetically distinct disorders. By combining patient-derived and isogenic hiPSC lines with longitudinal molecular, cellular, and ultrastructural analyses, we show that hOLS reproducibly generate a broad spectrum of CNS neuroglial cell types that undergo progressive maturation and display cellular properties and gene expression patterns that are relevant for studying globoid cell leukodystrophy (GLD), POLR3-related leukodystrophy (4H) and Canavan disease (CD). While this study primarily provides a framework for future studies addressing disease pathogenesis at higher scale and depth, our data indicate that disease-associated alterations emerge already during very early stages of human neural development, before extensive oligodendrocyte maturation and myelin formation, and involve multiple neural lineages beyond the oligodendrocyte compartment.

The hOLS system builds on myelinating spheroid models previously established by the Pașca laboratory (Birey et al., 2017; Marton et al., 2019). Consistent with other ventralized spheroid protocols (Amin et al., 2024; James et al., 2021), hOLS generated diverse neuronal and glial populations and exhibited a developmental progression characterized by early neurogenesis followed by sustained gliogenesis and oligodendrocyte maturation, recapitulating key temporal features of human brain development (Braun et al., 2023; Emery & Lu, 2015; Rowitch & Kriegstein, 2010). Single-cell transcriptomic profiling substantially expanded the cellular resolution of this model, enabling the identification of 27 distinct cell populations, including a serotonergic neuronal population that was not previously detected using bulk approaches. Mapping to a developing human brain reference further revealed a so far undescribed broad regional representation with telencephalic, diencephalic, and midbrain identities contributing to the overall hOLS composition. In accordance with the patterning protocol used, these cell populations showed mostly ventral neural identities and covering mostly the first-trimester of human development (Braun et al., 2023; Quadrato et al., 2017). Together, these findings show that hOLS capture substantial cellular diversity and developmental progression across neuronal and glial lineages, providing the cellular context needed to investigate early leukodystrophy-associated alterations.

One important finding of this study is that disease-associated alterations emerged before extensive oligodendrocyte maturation and myelin formation. Across all three leukodystrophies, transcriptional abnormalities were already evident in progenitor, immature neuronal, and glial populations by D100, when hOLS still contained substantial fractions of progenitor and immature neural cells. These early changes affected multiple lineages, including radial glia, glial progenitors, neuronal progenitors, and immature neurons, suggesting that pathogenic processes already begin during neural development rather than arising solely from defects in mature oligodendrocytes or myelin maintenance. Our observations therefore support the emerging view that leukodystrophies can involve broader neurodevelopmental abnormalities affecting progenitor states, cellular maturation, and tissue homeostasis (Ceravolo et al., 2024; Ferrer et al., 2026; Mangiameli et al., 2021; Nowacki et al., 2022). The presence of such alterations before extensive myelination further suggests that understanding the earliest stages of disease pathogenesis may be important for defining appropriate windows and cellular targets for therapeutic intervention.

The recurrent involvement of radial glia was particularly notable. Across the three LKDs, radial glial populations showed prominent but disease-specific transcriptional alterations, involving developmental programs in GLD, neurogenic differentiation programs in 4H, and metabolic pathways in CD. In 4H, oRG cells in particular showed strong enrichment of neurodevelopmental programs that were oppositely regulated in other progenitor and neuronal populations, pointing to altered coordination of developmental programs across neural lineages. As radial glia, including outer radial glia, serve as developmental progenitors for neuronal and glial lineages (Götz & Huttner, 2005; Hansen et al., 2010; Kriegstein & Alvarez-Buylla, 2009), their perturbation may represent an upstream point of vulnerability with consequence for both neurogenesis and gliogenesis. Recent studies have further shown that medial ganglionic eminence radial glia cells act as long-term source of interneurons in the developing human brain (Jia et al., 2026), raising the possibility that perturbations of ventral radial glia may contribute to the alterations in inhibitory neuronal populations observed in our hOLS. Developmental abnormalities involving neural progenitors have been previously reported in models of GLD, 4H and CD (Chapleau et al., 2025; Grønbæk-Thygesen & Hartmann-Petersen, 2024; Mangiameli et al., 2021; Perrier et al., 2021), highlighting early neural progenitors as potential therapeutic targets (Feng et al., 2020; Goldman et al., 2021). Together, these observations suggest that perturbation of early progenitor states in LKDs may contribute to downstream abnormalities across multiple neural lineages.

Despite their distinct genetic etiologies, the leukodystrophies examined here converged on alterations in glial metabolic and stress-response pathways. Astrocytic populations were prominently affected across all three disorders, supporting increasing evidence that leukodystrophy pathology extends beyond oligodendrocytes and involves broader glial dysfunction (Mangiameli et al., 2021; Nowacki et al., 2022). The underlying transcriptional programs however differed substantially between diseases. In GLD, astrocytic and oligodendroglial populations showed alterations in lysosomal, autophagic, mitochondrial and translational programs, consistent with the consequences of GALC deficiency and lysosomal dysfunction. In 4H, astrocytic populations displayed alterations in mitochondrial, ciliary, and microtubule-associated pathways, extending the consequences of POLR3 dysfunction beyond the oligodendrocyte lineage. In CD, astrocytes and immature astrocytes showed particularly prominent metabolic alterations, including changes in oxidative phosphorylation, mitochondrial function, translation, and chromatin-associated programs. These findings support an emerging view of leukodystrophies as multicellular disorders in which astrocytic and other glial populations are affected alongside oligodendrocyte lineage cells (Ceravolo et al., 2024; Nowacki et al., 2022; van der Knaap & Bugiani, 2017). Importantly, the distinct direction and magnitude of these responses across diseases and cell types argue against a uniform glial stress response and instead point to disease- and lineage-specific responses to different primary molecular defects.

In GLD, hOLS recapitulated the defining biochemical hallmarks of the disease, namely loss of GALC activity and psychosine accumulation (Feltri et al., 2021; Suzuki & Suzuki, 1970). GALC and ASAH1 were expressed across multiple neural populations, with particularly enrichment of ASAH1 expression within glial progenitor populations. This suggests that psychosine generation may begin during early gliogenesis rather than exclusively within mature oligodendrocytes, consistent with evidence for fetal-onset pathology in GLD (Feltri et al., 2021; Inamura et al., 2018; Kobayashi et al., 1988). Beyond these canonical biochemical features, showed widespread alterations in lysosomal, autophagic, mitochondrial, and translational pathways across progenitor, neuronal, and glial populations, supporting metabolic vulnerability as a central component of GLD pathogenesis (Mangiameli et al., 2021; Voccoli et al., 2014; Won et al., 2016). Importantly, in contrast to previous hiPSC-derived 2D cultures (Mangiameli et al., 2021, 2022), oligodendrocyte proportions in GLD hOLS remained relatively preserved despite profound biochemical and transcriptional abnormalities, suggesting that the multicellular 3D environment captures dysfunction preceding overt lineage depletion. This interpretation aligns with studies in Twitcher mice, which showed neuronal, axonal, and glial abnormalities before extensive demyelination (Potter et al., 2013; Taniike & Suzuki, 1994; Teixeira et al., 2014). Together, our data suggest that GALC deficiency induces early, lineage-specific cellular stress responses that precede extensive myelination and extend beyond mature oligodendrocytes.

In 4H hOLS, our findings extend the consequences of POLR3 dysfunction beyond the oligodendrocyte lineage and point to early alterations in progenitor and neuronal populations. The broad expression of POLR3A and POLR3B was consistent with their fundamental role in RNA polymerase III in cellular homeostasis, (Bernard & Vanderver, 1993; Choquet et al., 2019; Kok et al., 2025), while reduced POLR3A transcript levels were observed in only one of the two patient lines, suggesting that altered steady-state transcript abundance is not a general feature of the disease. Instead, pathway analyses revealed widespread alterations in RNA metabolism, translation, mitochondrial function, and neurodevelopmental programs across multiple lineages. The strongest transcriptional abnormalities occurred in progenitor and interneuron-associated populations, with neurodevelopmental programs showing opposing enrichment patterns between oRG, neuroepithelial cells, immature neurons, and SST neurons. Alterations affecting interneuron-related populations were consistent with previous evidence implicating abnormal interneuron development and neuronal dysfunction in POLR3-related leukodystrophy (Chapleau et al., 2025; Dooves et al., 2023; Perrier et al., 2021). Alterations in Hedgehog-related pathways further support a potential disturbance of developmental signaling in 4H, although the heterogeneous and cell type-specific responses observed here suggest that the consequences of POLR3 dysfunction differ substantially across neural lineages. Together, our data indicate that 4H involves early and widespread disruption of developmental and RNA-associated programs, providing a potential cellular basis for the combined neuronal and glial abnormalities characteristic of POLR3-related leukodystrophy.

In CD hOLS, ASPA deficiency and the expected accumulation of N-acetylaspartate (NAA), recapitulated a core metabolic feature of the disease (Baslow, 2002; Hoshino & Kubota, 2014; Matalon et al., 1988). Beyond this, our results revealed prominent astrocyte-centered metabolic dysfunction in accordance with previous evidence implicating astrocytes in CD pathogenesis (Grønbæk-Thygesen & Hartmann-Petersen, 2024; Namboodiri et al., 2006). Specifically, single-cell analysis showed dysregulation of mitochondrial and oxidative phosphorylation pathways in astrocytes, consistent with their proposed role as key metabolic mediators for energy supply and NAA/NAAG catabolism in white matter (Baslow & Guilfoyle, 2009). In oligodendroglial lineage cells, we noted reduced expression of multiple oxidative phosphorylation complexes, consistent with reports of oligodendrocyte-specific energetic deficits in CD, supporting a broader impact of ASPA deficiency on energy homeostasis (Francis et al., 2012, 2016; Madhavarao et al., 2005). Importantly, metabolic pathway alterations were also detected in several neuronal populations, despite the lack of ASPA expression in neurons, suggesting that ASPA deficiency exerts secondary non-cell-autonomous effects across neural lineages (Grønbæk-Thygesen & Hartmann-Petersen, 2024; Lotun et al., 2021; Namboodiri et al., 2006). We also observed gene expression signatures suggesting HAT- and HDAC-mediated dynamic alterations of acetylation across glial populations, which is in line with the notion that NAA catabolism impacts acetyl-CoA availability and histone acetylation, thus pointing to a possible effect of ASPA deficiency on epigenetic regulation and metabolic pathways (Dominicis et al., 2023; Lotun et al., 2023; Prokesch et al., 2016). Together our findings support a prominent role for astroglial metabolic dysfunction in CD while also revealing broader effects across oligodendrocyte-lineage and neuronal populations.

An important strength of this study is the integration of multiple leukodystrophies within a single experimental framework. By applying the same differentiation protocol, sampling strategy, and analytical pipeline across GLD, 4H, and CD hOLS, we were able to directly compare disease-associated alterations while minimizing technical variability. The hOLS model contains co-developing neuronal, astrocytic, and oligodendroglial populations and therefore enables disease-associated alterations to be examined across interacting neural lineages within the same 3D system. Moreover, the longitudinal design, combined with bulk and single-cell transcriptomics, molecular and metabolic analyses, and ultrastructural characterization, allowed us to distinguish alterations emerging at earlier developmental stages from those associated with later cellular maturation. Divergent results between bulk and single-cell datasets, particularly for lysosomal and inflammatory pathways in GLD, likely reflect opposing responses among distinct cellular populations that become obscured when averaged across heterogeneous tissues (Hwang et al., 2018; Lake et al., 2018). The inclusion of both patient-derived and gene-edited isogenic lines further enabled the study of disease phenotypes across different genetic backgrounds while providing genetically matched controls for selected disease models. Importantly, applying the same differentiation and analytical framework to GLD, 4H, and CD allowed direct comparison of disorders with distinct primary molecular defects and revealed both convergent developmental and glial alterations and disease- and lineage-specific transcriptional responses. This multiscale approach therefore provides a more comprehensive view of leukodystrophy pathogenesis, linking global transcriptional changes to the cellular populations in which they arise (Braun et al., 2023; Crowell et al., 2020).

Several limitations should nevertheless be considered. First, although hOLS reproducibly generated diverse neural populations, regional specification remained biased toward ventral forebrain and midbrain identities, potentially limiting the modeling of region-specific pathology (Amin et al., 2024; Marton & Pașca, 2020). Second, oligodendrocyte maturation did not fully reach postnatal states, and compact myelin formation remained incomplete, restricting analyses of later disease stages, a limitation shared by many current myelinating organoid systems (James et al., 2021; Kim et al., 2019; Madhavan et al., 2018; Marton & Pașca, 2020). As with other organoid-based systems, variability in cellular composition and maturation persisted between spheroids, reflecting the intrinsic self-organizing properties of 3D neural cultures (Scuderi et al., 2025; Velasco et al., 2019). Third, microglia, vasculature, peripheral immune components, and systemic influences were absent, constraining investigation of inflammatory and non-cell-autonomous mechanisms that likely contribute to disease progression *in vivo*, as previously highlighted for brain organoid models (Cakir et al., 2019; Kistemaker et al., 2025; Ormel et al., 2018). Fourth, the relatively small number of patient and isogenic lines also limits the ability to capture the full spectrum of disease heterogeneity. Future studies incorporating larger cohorts and additional disease-causing variants will be important to validate the generalizability of the mechanisms identified here. Finally, pathway enrichment analyses identify transcriptional programs associated with disease but do not by themselves establish functional pathway impairment or causal relationships between altered cell populations. Functional perturbation studies and larger cohorts will therefore be required to determine which of the lineage-specific signatures identified here directly contribute to disease pathogenesis.

In summary, this study establishes hOLS as a human 3D model for investigating the pathogenesis of leukodystrophy across developmental stages and disease subtypes. By comparing three genetically distinct leukodystrophies within a common experimental framework, we show that disease-associated changes emerge before extensive myelination and extend beyond the oligodendrocyte lineage to involve progenitor, neuronal, and astroglial populations. While some common disruptions in developmental and homeostatic pathways were observed in genetically distinct leukodystrophies, each disorder retains characteristic lineage-specific molecular signatures that reflect its underlying genetic defect. Beyond providing insight into GLD, 4H, and CD-associated alteration in early neural development, the dataset generated here constitutes a resource for use and further extension by the broader leukodystrophy community and thus a framework for future mechanistic studies, therapeutic target identification, and evaluation of emerging treatment strategies in a human developmental context.

## Methods

### hiPSC editing and culture

hiPSCs were generated at the respective institutes according to previously described protocols (Dooves et al., 2023; Holmes & Heine, 2017; Mangiameli et al., 2021; Rehbach et al., 2019). All subjects have given informed consent in accordance with the Declaration of Helsinki. Briefly, fibroblasts were reprogrammed into iPSCs using either the CytoTune™-iPS 2.0 Sendai Reprogramming Kit (Invitrogen), polycistronic construct with *OCT4*, *SOX2*, *KLF4*, and *C-MYC* within a lentiviral vector or StemMACS mRNA Reprogramming Kit (Miltenyi Biotec). Pluripotency was confirmed by immunocytochemistry, alkaline phosphatase assay, PCR, embryoid body formation, pluritest. Karyotype analysis was also performed to exclude chromosomal abnormalities.

GALC KO and KI hiPSCs were produced by CRISPR/Cas9-mediated gene editing of the HD2 hiPSC line. GALC KO hiPSCs were generated using the Gene Knockout Kit v2 (Synthego), designed to target the first exon of the human GALC gene, according to the manufacturer’s instructions. Generation of GALC KI hiPSC was outsourced to Synthego and was obtained by inserting the pathogenic G553R mutations (*GALC*, c.1657G>A) in both alleles.

Following generation, gene editing, and characterization, all hiPSC culture procedures were centralized at the IRCCS San Raffaele Scientific Institute. hiPSCs originating from the IRCCS San Raffaele Scientific Institute and the Institute of Reconstructive Neurobiology were thawed and maintained on Matrigel (Merck Millipore)-coated plates in StemMACS iPS-Brew XF medium (Miltenyi Biotec). Medium was refreshed daily, and cells were maintained at 37°C in a humidified atmosphere containing 5% CO₂ and 5% O₂. Upon reaching confluence, cells were passaged at ratios ranging from 1:3 to 1:10 using 0.5 mM EDTA (Thermo Fisher Scientific) diluted in PBS.

hiPSCs originating from the Vrije Universiteit were initially thawed under the culture conditions used at the institute of origin, using TeSR™-E8™ medium (STEMCELL Technologies) supplemented with 10 µM ROCK inhibitor Y-27632 (Selleckchem) on Vitronectin XF™-coated plates (STEMCELL Technologies). At the subsequent passage, cells were transferred to Matrigel-coated plates and adapted to iPS-Brew medium. An overview of the hiPSC lines used in this study is provided in Table 1.

To generate ASPA KO and isogenic control lines for CD, hiPSCs were cultivated in iPS-Brew XF medium (Miltenyi) on vitronectin until they were 70% confluent. 1 hour before nucleofection, the medium was changed to iPS-Brew XF supplemented with 10 µM ROCK Inhibitor Y-27632 (BioTechne,). For nucleofection, hiPSCs were dissociated using StemPro Accutase (A11105) supplemented with 10 µM Y-27632 and 6 x 10^5^ cells were resuspended in 20 µL nucleofection solution (P3 Primary Nucleofection Kit, Lonza) containing 2 µM Alt-R Cas9 Electroporation Enhancer (IDT) and a CRISPR-Cas9 ribonucleoprotein (RNP) complex composed of 4 µM Alt-R HiFi Cas9 Nuclease (IDT) and a 5 µM tracrRNA:crRNA duplex (spacer sequence: TCCGGTTAGCTCATTCCCAT). The cells were then subjected to program CM150 of the Amaxa 4D Nucleofector (Lonza) and nucleofected cultures were expanded for subcloning. For CD, three isogenic hiPSC lines (CD 1i, CD 2i, CD 3i), each harboring frameshift mutations in the ASPA gene (exon 1) were genotyped using the Phire Animal Tissue Direct PCR Kit (Thermo Scientific) and subsequent Sanger sequencing (Microsynth Seqlab).

### hOLS generation

Spheroids from a total of five control, three Globoid Leukodystrophy, three Canavan disease, and two 4H leukodystrophy hiPSC lines (See Table 1) were generated according to an adapted version of Marton et al. (2019) protocol (Marton et al. 2019). Before differentiation, EB formation was started from single cell suspension after treatment with Accutase (Merck Millipor,) dissociation for 5-7 minutes. Cells were collected by centrifugation (5 min, 400 rcf) in an E8 medium (Thermo Fisher). Pellets were resuspended in E8 medium supplemented with ROCK inhibitor Y-27632 (10 µM, SIGMA). Uniform spheroids were obtained by dropwise plating of 3 x 10^6^ cells in each well of an AggreWell™800 (STEMCELL Technologies), which was pretreated with Anti-Adherence Rinsing Solution (STEMCELL Technologies). After plating, cells were captured in the bottom by centrifugation at 100 rcf for 3 minutes and placed in an incubator (37°C, 5% CO_2_). After 24 hours, spheroids of approximately 300 µm had formed in each of the 300 microwells. Spheroids were collected by pipetting up and down the medium with sterile wide-bore tips (Thermo Fisher) and transferred to low attachment 10 cm Petri dishes. To initiate the differentiation, on day 1 of the protocol, spheroids were plated in E6 medium (Thermo Fisher,) supplemented with SMAD pathway inhibitors dorsomorphin (2.5 µM, Sigma-Aldrichand SB-431542 (10 µM, Sigma-Aldrich). The medium was refreshed daily for the next two days. On days 4 and 5 also, Wnt pathway inhibitor IWP-2 (5 µM, Selleckchem) was used in the medium. The following six days medium was still refreshed daily but with Differentiation and Maintenance Medium (DMM), which consisted of DMEM/F12 1:1 (Gibco) (Thermo Fisher Scientific) with 1x B-27 supplement without vitamin A (Thermo Fisher Scientific,), 1x N2 supplement (Invitrogen), 1x MEM NEAA (Invitrogen), 1x Glutamax (Invitrogen), 25 µg/ml Human Insulin (Sigma-Aldrich), 0.1 mM β-mercaptoethanol (Invitrogen), 100 U/ml P/S and supplemented with 20 ng/ml EGF (Peprotech) 20 ng/ml FGF-2 (Peprotech) and five µM IWP-2 (Selleckchem). Between days 12 and 17, 1 µM of Smoothened agonist (Cayman) was supplemented to the daily refreshed medium. From day 18 onward, the frequency of medium changes was reduced to every other day. On day 25, DMM was supplemented with 60 ng/ml T3 (Sigma-Aldrich), 100 ng/ml Biotin (Sigma-Aldrich), 20 ng/ml NT-3 (Peprotech), 20 ng/ml BDNF (Peprotech), 1 μM cAMP (Sigma-Aldrich), 5 ng/ml HGF (Peprotech), 10 ng/ml IGF-1 (VWR), 10 ng/ml PDGF-AA (R&D Systems). On day 37, DMM was supplemented with 60 ng/ml T3, 100 ng/ml Biotin, 1 μM cAMP, and 20 μg/ml Ascorbic Acid (Sigma-Aldrich). Spheroids were maintained in this medium until the end of the experiment. From day 44 onwards, the medium changes were reduced to twice weekly.

### hOLS embedding and sectioning

hOLS were collected for embedding on the following days of differentiation (D) 50, D100, and D150. After fixation in 4% PFA for 15 minutes, and a thorough PBS wash, spheroids were allowed to dehydrate overnight in 30% sucrose (in PBS) at 4°C. Then, they were transferred to embedding molds with OCT embedding medium (Bio-Optica) and snap-frozen in an isopropanol bath. Frozen samples were stored at -80°C until cryosectioning. Cryosections of 20 µm were made using a standard cryostat and transferred to Superfrost® Plus adhesive microscope slides (Thermo Scientific). hOLS slices were stored at -80°C until immunofluorescence staining.

### Immunofluorescence

Cryosectioned hOLS were rehydrated with PBS to perform immunofluorescence and incubated for 1 hour with blocking solution (PBS containing 0.3% TritonX-100 (Sigma-Aldrich), and 10% Normal Goat Serum (NGS, Sigma-Aldrich). Primary antibodies diluted in blocking solution were incubated overnight at 4°C (See Supplementary Table 1). The following day, slides were washed 3 times with PBS and incubated for 2 hours at room temperature with secondary antibodies (Supplementary Table 2) diluted in PBS with 1% NGS. After incubation, the slides were washed with PBS 3 times. Nuclear counterstaining was performed with Hoechst (1:1000, in PBS) for 10 minutes at room temperature. A final wash with PBS was performed before mounting coverslips using FluorSave mounting medium (VWR). Images were acquired at the Advanced Light and Electron Microscopy Bioimaging Center (ALEMBIC) at IRCCS San Raffaele Scientific Institute using a Mavig RS-G4 confocal microscope and a 40x oil immersion objective. FiJi and Imaris software (NIH, Bethesda, MD) were used for image visualization and analysis.

### Electron microscopy

hOLS were fixed using 2.5% glutaraldehyde in 0.1 M cacodylate buffer (pH 7.4) overnight at 4°C. Samples were post-fixed for 1 hour in a reduced osmium solution (1.5% potassium ferricyanide and 1% aqueous osmium tetroxide in 0.1 M cacodylate buffer). Following the initial incubation with the heavy metal-based solutions, the pellets were washed with bi-distilled water at room temperature, immersed in a 0.5% uranyl acetate solution, and left overnight at 4 °C. The samples were dehydrated using a graded ethanol series (70%, 80%, 90%, 100%) and acetone for 10 minutes each before being embedded in Epon resin. After curing at 60 °C for 48 hours, thin sections were cut using a Leica UC7 ultramicrotome (Leica Microsystems). The sections were mounted onto 300-mesh copper grids and imaged using a Talos L120C G2 transmission electron microscope (Thermo Fisher Scientific) at an acceleration voltage of 120 kV at the Advanced Light and Electron Microscopy Bio-Imaging Centre of San Raffaele Scientific Institute (ALEMBIC).

### Bulk mRNA sequencing

According to the manufacturer’s protocol, total RNA was extracted from individual spheroids with the RNeasy Mini Kit (Qiagen). Per donor, RNA from three spheroids was pooled to create a single bulk mRNA sequencing sample. The RNA concentration and purity were determined using a NanoDrop spectrophotometer (Thermo Fisher Scientific). Purified RNA samples were sent to Genewiz (Azenta Life Sciences) for RNA sequencing (RNA-seq) analysis. The library was prepared with the NEBnext Ultra II Directional RNA Library Preparation kit (New England Biolabs), and sequencing was performed on an Illumina NovaSeq 6000 platform. A minimum read depth of 40 million per sample was achieved.

Following quality control with FastQC, preprocessing was performed using TrimGalore (v0.5.0) to remove adapters and low-quality bases. The trimmed reads were then aligned to the GRCh38 reference genome assembly provided by the 10X reference data repository (refdata-cellranger-GRCh38-3.0.0) using STAR (v2.7.0d). The gene annotation file provided by 10X Genomics (refdata-cellranger-GRCh38-3.0.0.gtf), which includes exon and transcript intervals, was used to ensure optimal compatibility with scRNA-seq data. Post-alignment quality metrics—such as coverage distribution across gene bodies and the percentage of reads mapping to exons—were assessed using QoRTs (v1.3.6). Read assignment to genes was performed with featureCounts (v1.6.3), using a minimum overlap of 10 bases and excluding chimeric reads.

Count data were analysed with the DESeq2 R package (v1.26.0), without prefiltering of low-count genes. For the exploratory analysis shown in Fig. 4A, counts were transformed with the regularised logarithm (rlog) and principal component analysis was performed on the transformed values. Differential expression was assessed separately at D50, D100 and D150, using the design ∼ batch + condition, where batch corresponds to the institute of origin of the hiPSC lines, with HD as the reference level. P-values were adjusted with the Benjamini–Hochberg procedure, and genes were ranked by log2 fold change for gene set enrichment analysis.

### Single-cell RNA-seq (scRNA-seq)

To obtain a single cell suspension from 3D brain spheroids, 3 spheroids from the same genotype and batch at D100 or D150 were pooled and gently dissociated to obtain single-cell transcriptomic data. Brain spheroids were harvested from culture dishes and mechanically dissociated by mincing them into smaller pieces using a sterile scalpel blade (Bisturi) for approximately 2–3 minutes until spheroids were visibly fragmented into fragments smaller than 1 mm in diameter. Care was taken to avoid excessive mechanical force and minimize cell damage. Following mechanical trituration, enzymatic dissociation was performed using the Papain Dissociation System (Worthington Biochemical Corporation) following the manufacturer’s instructions, with minor modifications. Briefly, the minced spheroids were incubated with papain and transferred to an orbital shaker at 70 rpm inside a tissue culture incubator for 70 minutes. Dissociated fragments were pelleted and resuspended with a papain inhibitor to stop the reaction. After resuspending in 0.2 % bovine serum albumin diluted in PBS and supplemented with 10uM Y-27632, cells were filtered through a 70 µm cell strainer to remove undissociated tissue clumps. Cell counting was performed using TC20 automated cell counter (BioRad), and viability was calculated as the percentage of live cells (trypan blue-negative) over the total cell count. Each sample’s viability was evaluated to ensure it was above 90%. Finally, cells were suspended in sterile PBS 0.04% BSA to a final concentration of 1 × 10^6^ cells/mL (1,000 cells/μL).

Single cells were processed using the Chromium Single Cell 3’ Reagent Kit (v3.1 Chemistry) and the 10x Chromium Controller platform (10X Genomics), according to manufacturer’s instructions, targeting the recovery of 4000 cells. The resulting libraries were evaluated for size distribution and concentration using HS DNA assay on Tape Station 4200 (Agilent Technologies)

Libraries were sequenced on Illumina NovaSeq6000 to obtain approximately 50.000 reads per cell. Single-cell encapsulation, cDNA amplification, library prep, and sequencing were performed at COSR (Center for Omic Sciences, San Raffaele Hospital). The Cell Ranger software suite (version 6.1.1, 10X Genomics) was used to demultiplex the data, align reads to the human reference genome (GRCh38) and quantify Unique Molecular Identifier (UMI). From 25 samples, we obtained a total of 144,087 cells, with a median of 49,018 mean reads per cell and a median of 8945 UMI counts per cell. Raw counts were processed in Seurat (v4.1.1). Low quality cells were filtered out based on mitochondrial content higher than 20%, and cells expressing fewer than 200 or more than 6,000 genes were excluded. Log-normalization of UMI counts were performed with a scale factor of 10,000. The top 20% of variable genes were selected for the downstream analysis. Cell cycle scores were calculated using the CellCycleScoring function and a reference gene lists (Braun et al., 2023). The cell cycle difference was defined as the difference between S phase and G2/M phase scores.

Data were scaled, and unwanted sources of variability, such as UMI count, percentage of mitochondrial genes (cells above 20% were excluded) and cell cycle difference, were regressed out. Downstream analysis was performed on the top 30 principal components (PCs) out of 50. The Harmony package (v1.0) (Korsunsky et al., 2019) was used to mitigate patient- and site-related biases. Clustering resolution of 0.6 was used and marker genes computed using the FindAllMarkers function, which exploits a Wilcoxon Rank Sum test for significance, with logFC threshold set at 0.25. P-values <1e−5 were used to address clusters. Cell atlas of the first-trimester developing human brain (Braun et al., 2023), PanglaoDB and customized manually curated gene signatures were used as reference for cluster annotation on HD spheroids. To annotate the LKD samples, the HD dataset was used as a reference for anchor-based label transfer. We further explored spheroid regionalization by transferring cell labels from a cell atlas of the first-trimester developing human brain (Braun et al., 2023). This atlas comprises samples collected at multiple time points spanning 5 to 14 post-conception weeks (pcw). Anchors between the reference atlas and the hOLS query datasets were identified with the FindTransferAnchors function, and cell-type labels were assigned with TransferData (Seurat). To assess how regional identity assignments depend on the developmental stage of the reference, label transfer was performed (i) against the full atlas (all available time points, 5–14 pcw) and (ii) against two developmental windows analyzed separately, 5–7 pcw and 8–14 pcw.

Disease effects on cell type composition were calculated by transforming cell type counts using the centered log-ratio (CLR) transformation (Aitchison, 2003), and modeled independently for each cell type as CLR ∼ disease + time + disease × time (OLS, with HD and D100 as reference); disease effects at each timepoint were tested using the model coefficients and corrected for multiple testing using the Benjamini-Hochberg FDR procedure (* < 0.1, ** < 0.05, *** < 0.01).

Differential expression analysis between each leukodystrophy and HD hOLS was performed separately for each annotated cell type at D100 and D150, using the FindMarkers function of Seurat on log-normalised values of the RNA assay. Cells were grouped by disease condition within each cluster, with HD as the reference group, and significance was assessed with a Wilcoxon rank-sum test. Genes were ranked by average log2 fold change and the resulting lists were used as input for gene set enrichment analysis.Gene Set Enrichment Analysis (GSEA) was performed for each cell type within each comparison on the Log2FC-ranked gene lists, using the GSEA function of the clusterProfiler R package (v4.8.1), against the following MSigDB collections (v2023.1.Hs): Hallmark (H), the KEGG and Reactome canonical pathway subsets (C2: CP:KEGG and CP:REACTOME), and Gene Ontology (C5: GO:BP, GO:CC and GO:MF). Enrichment significance was estimated using the adaptive multilevel permutation procedure implemented in fgsea (v1.26.0). Normalized enrichment scores (NES) and Benjamini–Hochberg-adjusted p-values were calculated for all tested gene sets. The R package ggplot2 (v3.4.2) was used to graphically represent NES. HALLMARK terms were ranked by the sum of their NES, and terms with adjusted p-values below 0.05, 0.01 and 0.001 were highlighted. The same procedure, collections and thresholds were applied to the ranked gene lists obtained from the bulk RNA-seq comparisons.

Trajectory inference was performed with the R package Slingshot (v2.2.1) (Street et al., 2018), using cluster labels as input and the UMAP embedding computed on the Harmony-corrected principal components as the reduced-dimensional space. Trajectories were inferred on the whole dataset, including all genotypes and both collection time points, so that a single pseudotime axis was defined for all cells, with the NE cluster set as the starting cluster. End clusters were not specified, and terminal states were therefore inferred from the data, while all other parameters were left at their default values. The resulting lineages describe the progressive transition from neuroepithelial cells to oligodendrocytes, astrocytes and neurons, and pseudotime values were extracted for each cell along each lineage.

Pseudotime distributions were then compared between the two time points for each lineage, restricted to healthy donor samples and to cells with a lineage assignment weight above 0.5. Comparisons were performed across the five healthy donors, using the median pseudotime per donor at each time point and a paired Wilcoxon signed-rank test, with Benjamini–Hochberg correction across lineages. All donor–lineage–time point combinations were supported by at least 20 cells. Pseudotime was also compared between healthy and leukodystrophy hOLS, separately for each lineage and time point, using linear mixed models with genotype as fixed effect and cell line as random effect, fitted on cells with a lineage assignment weight above 0.5. P-values were adjusted with the Benjamini–Hochberg procedure across all comparisons.

### GALC activity assay

Specific beta-galactocerebrosidase activity was measured by hydrolysis of artificial fluorogenic substrate 4-methylumbelliferone-beta-galactopyranoside in presence of the competitive inhibitor of beta-galactosidase AgNO_3_, as previously described (Martino et al., 2009).

### (Lyso)glycosphingolipid measurement

Glucosylceramide (GlcCer), galactosylceramide (GalCer), glucosylsphingosine (LysoGlcCer), and galactosylsphingosine (LysoGalCer; psychosine) were quantified in GLD hiPSC and hOLS. The analytical procedure was adapted from previously described protocols for glycosphingolipid and lysosphingolipid quantification in plasma and serum (Groener et al., 2007; Sidhu et al., 2018; Voorink-Moret et al., 2018). Cell pellets were homogenized in 150 µL water by sonication on ice. A 50 µL aliquot was taken for determination of protein concentration using the bicinchoninic acid (BCA) assay. For lipid extraction, 75 µL of homogenate was mixed with internal standards consisting of 25 pmol LysoGlcCer-d5 (25 µL, 1 µM in methanol), 25 pmol LysoGalCer-d7 (25 µL, 1 µM in methanol), and 20 pmol GlcCer-d7 (10 µL, 2 µM in methanol). Subsequently, 240 µL methanol and 150 µL chloroform were added. After vortexing and incubation at room temperature, samples were centrifuged at 15,700 g for 10 minutes at 4°C to precipitate protein. The supernatant was transferred to a 2 mL tube and 150 µL chloroform and ammonium formate/formic acid buffer (225 µL, pH 3.15) were added. After vortexing, samples were centrifuged at 15,700 g for 3 minutes at 4°C to separate phases and the upper phase was dried under nitrogen at 40°C. After butanol/water partitioning, the upper phase was dried again and reconstituted in 100 µL of mobile phase B (95:5 acetonitrile:water with 1 mM ammonium formate and 0.1% formic acid). The lower phase (glycosphingolipids) was dried under nitrogen and deacylated by incubation with 0.5 mL of 0.1 M NaOH in methanol using a microwave program. After neutralization with 50 µL 0.1 M HCl in methanol, the processed samples were combined with the lysosphingolipid workflow from the drying step. UPLC-MS/MS analysis was performed on a Waters Acquity UPLC system coupled to a Waters Xevo TQ-XS mass spectrometer, operating in positive electrospray ionization (ESI) mode. Chromatographic separation was achieved on an Ascentis® Express HILIC column (4.6 × 50 mm, 2.7 μm, Supelco) with a HILIC Securityguard™ precolumn (4 × 3.0 mm, Phenomenex) at ambient temperature. The mobile phases consisted of (A) water with 1 mM ammonium formate and 0.1% formic acid, and (B) acetonitrile/water (95:5) with 1 mM ammonium formate and 0.1% formic acid. A gradient elution was applied at 1.5 mL/min with the following program: 0–0.2 min, 100–95% B; 0.2–3.5 min, 95% B; 3.5–4.0 min, 95–10% B; 4.0–5.0 min, 10% B; 5.0–5.1 min, 10–100% B; 5.1–7.0 min, 100% B. The injection volume was 10 µL. Analytes were detected using multiple reaction monitoring (MRM) of specific mass transitions. Quantification was performed against matrix-matched calibration curves using deuterated internal standards. Data acquisition and processing were performed using MassLynx software. Sphingolipid concentrations were normalized to the protein content of the homogenate and expressed as pmol per gram of protein for LysoGlcCer and LysoGalCer or nmol per gram of protein for GlcCer. Analyses were performed at Core Facility Metabolomics (CFM) Amsterdam UMC.

### Protein extraction and Western blot analysis

Spheroids (2-3 for each line) were washed twice with cold phosphate-buffered saline (PBS) and lysed in self-made RIPA buffer (50 mM Tris-HCl, pH 7.4; 150 mM NaCl; 1% NP-40; 0.5% sodium deoxycholate; 0.1% SDS) supplemented with protease and phosphatase inhibitor cocktails (Halt™ Protease and Phosphatase Inhibitor Cocktail (100X) Thermo Fisher Scientific). Lysates were incubated on ice for 60 minutes with intermittent vortexing (every 5’) and then centrifuged at 13,000 × g for 10 minutes at 4°C to remove cell debris. Supernatants were collected, and protein concentration was determined using the bicinchoninic acid (BCA) assay (Pierce™ BCA Protein Assay Kit, Thermo Fisher Scientific) following the manufacturer’s protocol.

Equal amounts of protein (20 µg) were mixed with 4× Laemmli sample buffer, boiled at 95°C for 5 minutes, and separated by SDS-PAGE on 15% polyacrylamide gels. Proteins were transferred onto PVDF membranes (Millipore) using a wet transfer system at 100 V for 90 minutes at 4°C. Membranes were blocked with 5% non-fat dry milk in Tris-buffered saline (TBS-T, 0.1% Tween-20) for 1 hour at room temperature and then incubated overnight at 4°C with primary anti-ASPA antibody (rabbit monoclonal; Abcam, ab223269)) diluted in blocking solution. After washing, membranes were incubated with HRP-conjugated secondary antibodies for 1 hour at room temperature. Protein bands were visualized using enhanced chemiluminescence (ECL) substrate (e.g., Bio-Rad Clarity or Thermo Fisher SuperSignal) and imaged with a chemiluminescence detection system (Immobilon^®^ Classico Western HRP substrate, Millipore/Merck). Target protein levels were normalized to loading controls such as TUBB3 (mouse monoclonal; Eurogentec). For detection of the human GALC protein, sample preparation, SDS–PAGE, and western blotting were performed as previously described (Mangiameli et al., 2021). After blocking, PVDF membranes were incubated with anti-GALC primary antibody (Abcam, ab137750) diluted 1:3,000 in 5% non-fat milk in TBST, followed by HRP-conjugated goat anti-rabbit secondary antibody (Millipore, 12-348) diluted 1:5,000. Immunodetection was performed using ClarityTM ECL Western Blotting Substrate (Biorad) and imaged with a ChemiDoc Imaging System (Biorad). Brightness and contrast of the blots were adjusted to enhance image quality without altering the informational content or misrepresenting the data.

### NAA analysis

N-acetyl aspartate (NAA) levels in the myelinoids was measured according to the enzymatic and fluorimetric method described in (Becker & Eckhardt, 2023).

### Statistics

Data were analyzed with GraphPad Prism version 8 for macOS and are expressed as mean ± standard error of the mean (SEM). The number of samples, the statistical tests and the post-tests used are specified in the figure legends. p < 0.05 was considered statistically significant. GSEA statistical significance was assessed using the adaptive multilevel permutation procedure implemented in fgsea, with Benjamini–Hochberg correction for multiple testing. Differential cell-type composition was assessed on centred log-ratio (CLR)-transformed counts using linear models, with Benjamini–Hochberg FDR correction.

## Acknowledgements

We thank Sabata Martino and Francesco Morena for quantification of GALC activity, and all the members of the Gritti lab for continuous support and helpful discussion.

Part of this work was carried out in the core facilities established at IRCSS San Raffaele Scientific Institute and Vita-Salute San Raffaele University: ALEMBIC (Advanced Light and Electron Microscopy BioImaging Center) and COSR (Center for Omics Sciences).

C.R. conducted this study as part of her PhD program in Molecular Medicine, curriculum of Gene and Cell Therapy, XXXIX cycle (Vita-Salute San Raffaele University).

## Funding

This study was supported by grants from: European Joint Programme on Rare Diseases (EJP RD JTC 2019 – NG4Leuko) to V.H; A.G; and O.B; Fondazione Telethon (TTAGD0222TT) to A.G.

## Notes

### Competing Interest Statement

The authors have declared no competing interest.

