## Supplemental for "Glia-enriched human brain organoids for studying developmental alterations in leukodystrophies"

### Supplementary Figure 1

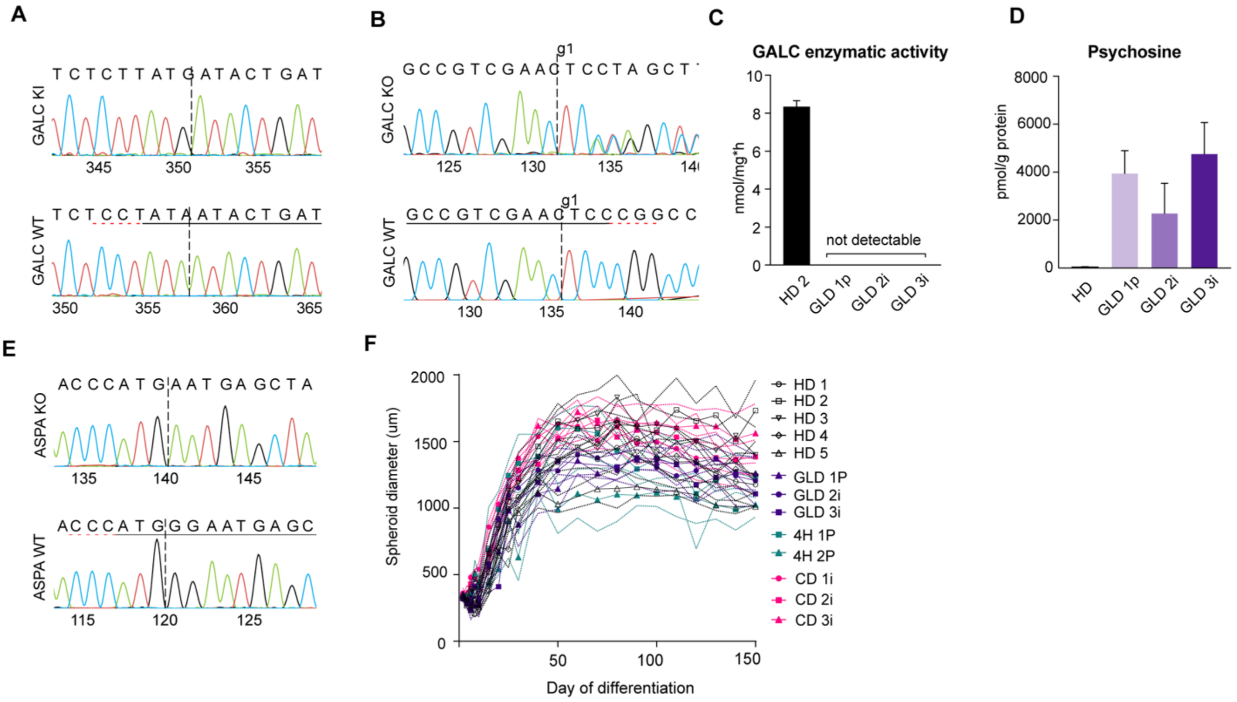


***Figure S1. Validation of GLD and CD hiPSC models and characterization of hOLS growth dynamics***

***(A)*** *Representative Sanger sequencing chromatograms of the GALC gene in HD and GALC knock-in (KI) hiPSCs. The wild-type sequence shows the expected nucleotide peaks, whereas the mutant sequence exhibits a biallelic 1657A>G transition in exon 14 of GALC, resulting in a G553R substitution. A silent C>T substitution at position 1653 is also present.* ***(B)*** *Representative Sanger sequencing chromatograms of the GALC gene in wild-type and GALC knockout (KO) hiPSC lines. The wild-type sequence shows the expected nucleotide peaks, whereas the mutant sequence exhibits a biallelic frameshift mutation in exon 2 of GALC.* ***(C)*** *GALC enzymatic activity in HD2 and GLD (GLD 1p, GLD 2i, GLD 3i) hiPSCs. Data are expressed as mean ± SEM (n = 2–3 replicates/clone).* ***(D)*** *Psychosine levels measured by mass spectrometry in HD (HD1, HD2) and GLD (GLD 1p, GLD 2i, GLD 3i) hiPSCs. Data are expressed as mean ± SEM (n = 2–6 replicates/clone).* ***(E)*** *Representative Sanger sequencing chromatograms of the ASPA gene in wild-type and ASPA-knockout hiPSCs. The wild-type sequence shows the expected nucleotide peaks, whereas the mutant sequence exhibits a frameshift mutation in exon 1 of ASPA.* ***(F)*** *Time-course measurements of hOLS diameter. Data are expressed as mean (symbols) ± SD (dotted lines) (n = 3–22 hOLS/group/time point).*

### Supplementary Figure 2

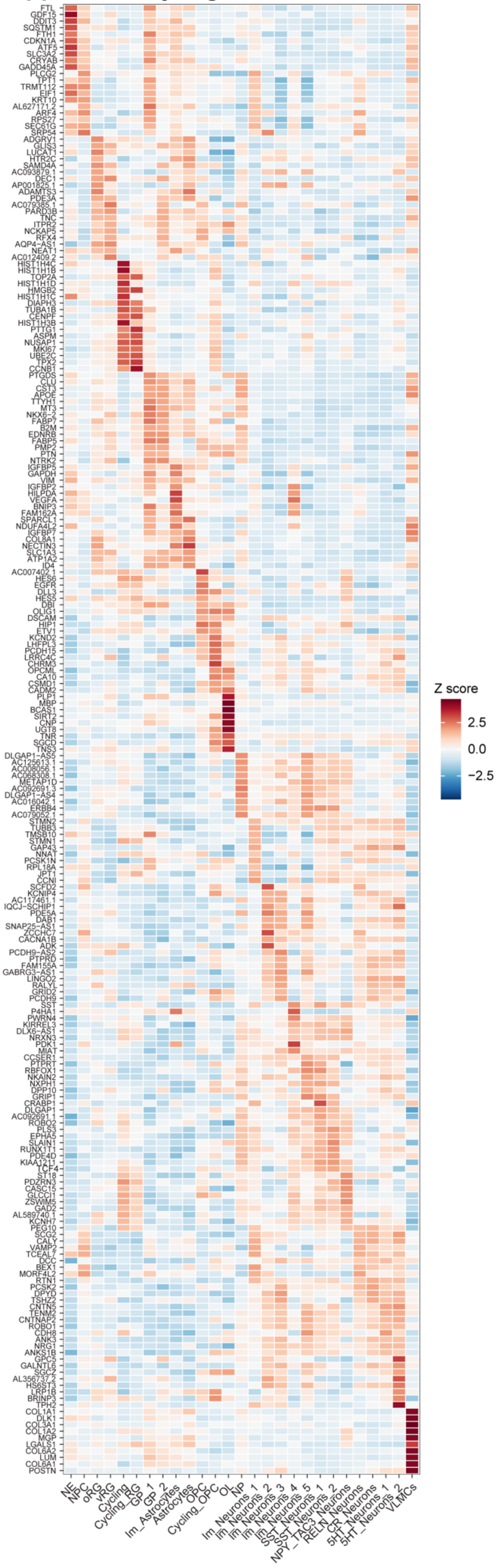


***Figure S2. Cluster-specific marker gene expression in HD hOLS***

*Heatmap showing the expression levels of the top 10 differentially expressed genes for each cell cluster identified by scRNA-seq analysis of HD hOLS. The color scale represents relative expression as Z-scores (blue, below average; red, above average). For expression of established cell-type marker genes, see Fig. 2B. For cell-type abbreviations, see the legend to Fig. 2.*

### Supplementary Figure 3

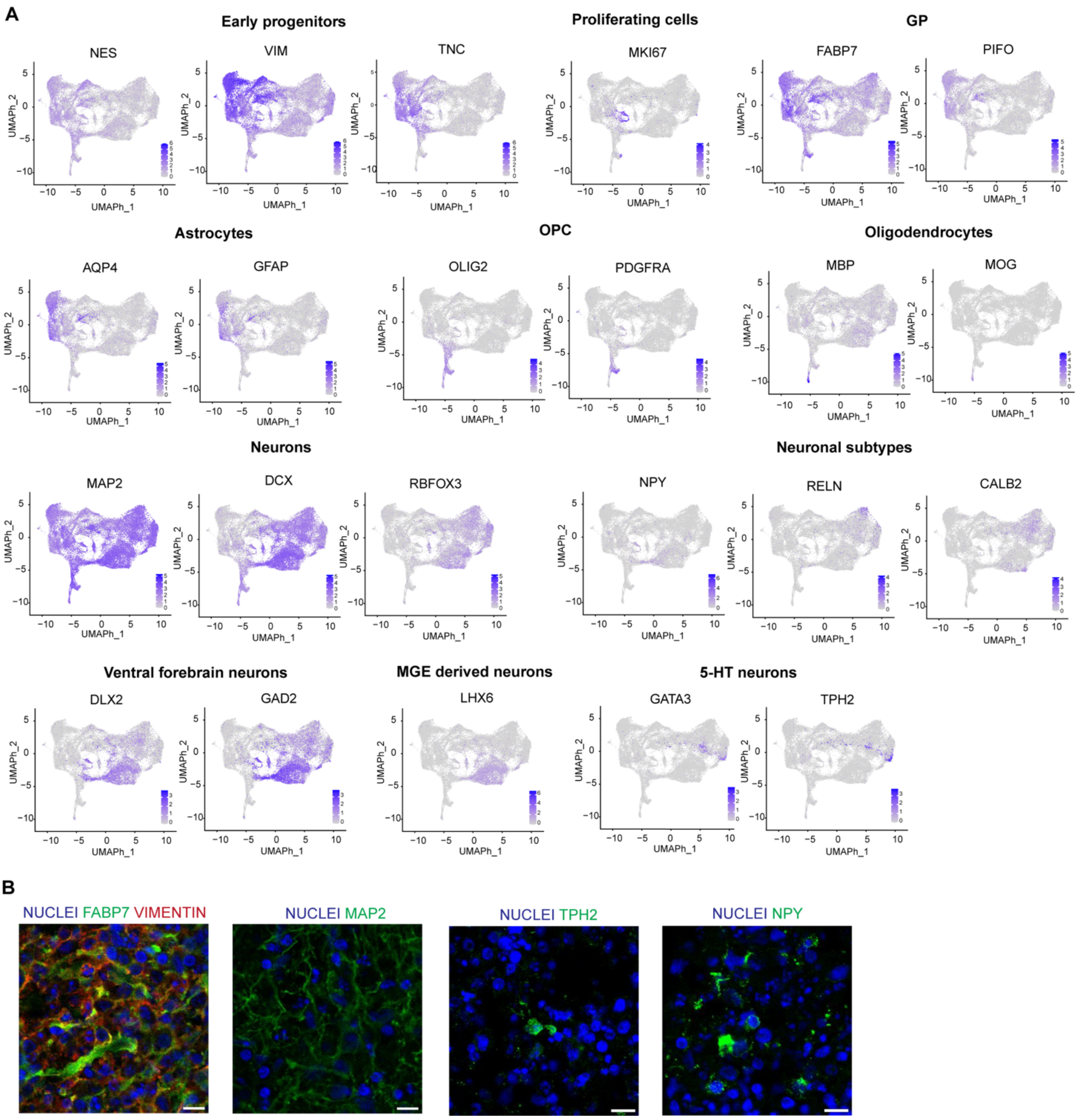


***Figure S3. Lineage-specific marker expression in HD hOLS***

*__(A)__ Feature plots showing expression of key lineage marker genes in HD hOLS projected onto UMAPs. Cells are colored according to gene expression level (grey, low; violet, high). __(B)__ Representative confocal images of immunofluorescence staining of hOLS sections at D150. From left to right: FABP7 (green), VIMENTIN (red), MAP2 (green), TPH2 (green), and NPY (green). Nuclei are counterstained with Hoechst (blue). Scale bar: 10 µm.*

*GP: Glial Progenitors; OPC: Oligodendrocytes Progenitors Cells; MGE derived neurons: Medial Ganglionic Eminence derived neurons; 5-HT neurons: Serotonergic neurons.*

### Supplementary Figure 4

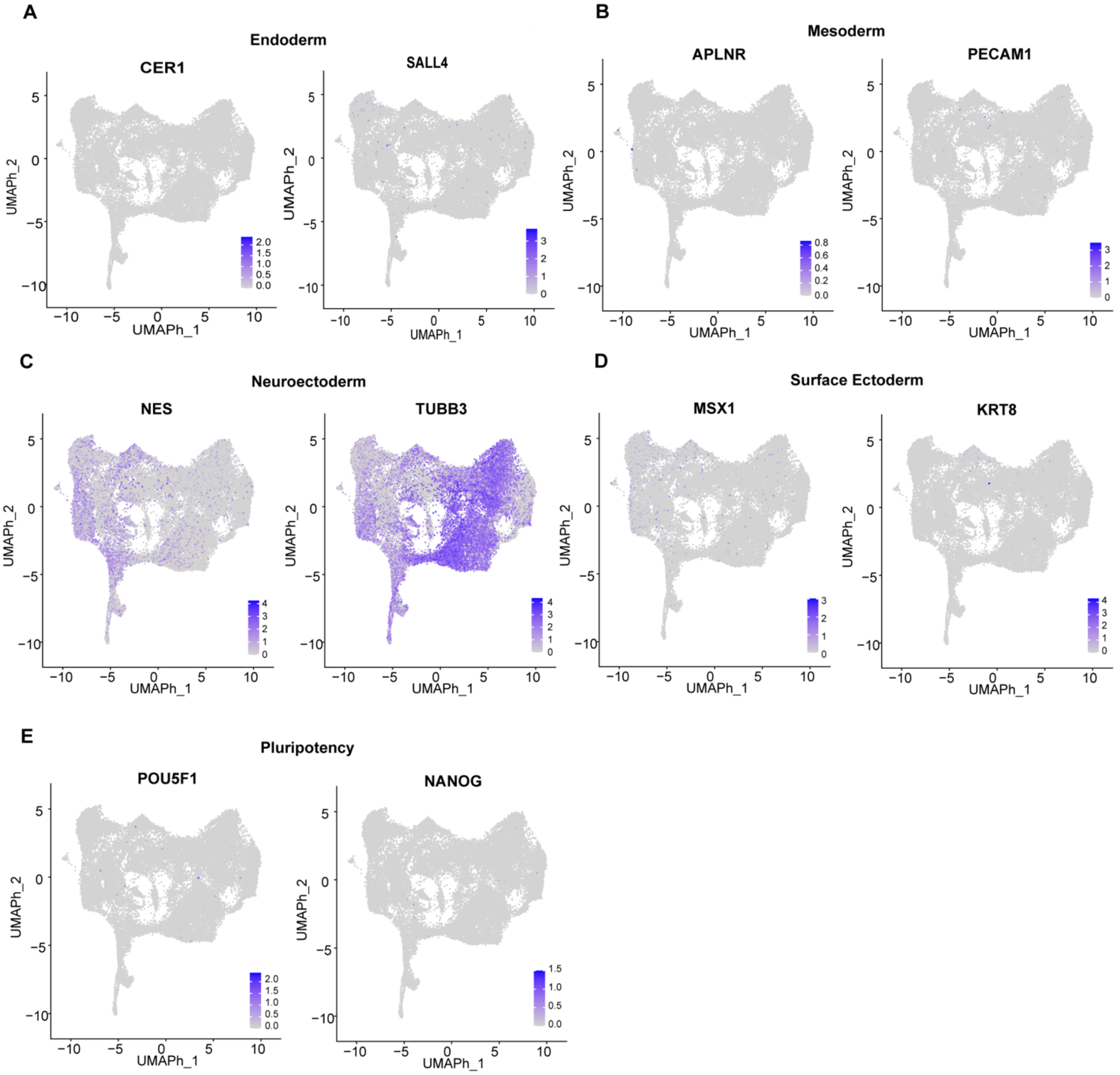


***Figure S4. Developmental identity of HD hOLS***

*Feature plots showing expression of genes associated with **(A)** endoderm, **(B)** mesoderm, **(C)** neuroectoderm, **(D)** surface ectoderm, and **(E)** pluripotency in HD hOLS. Expression levels are represented by the color gradient from grey (low) to blue (high).*

### Supplementary Figure 5

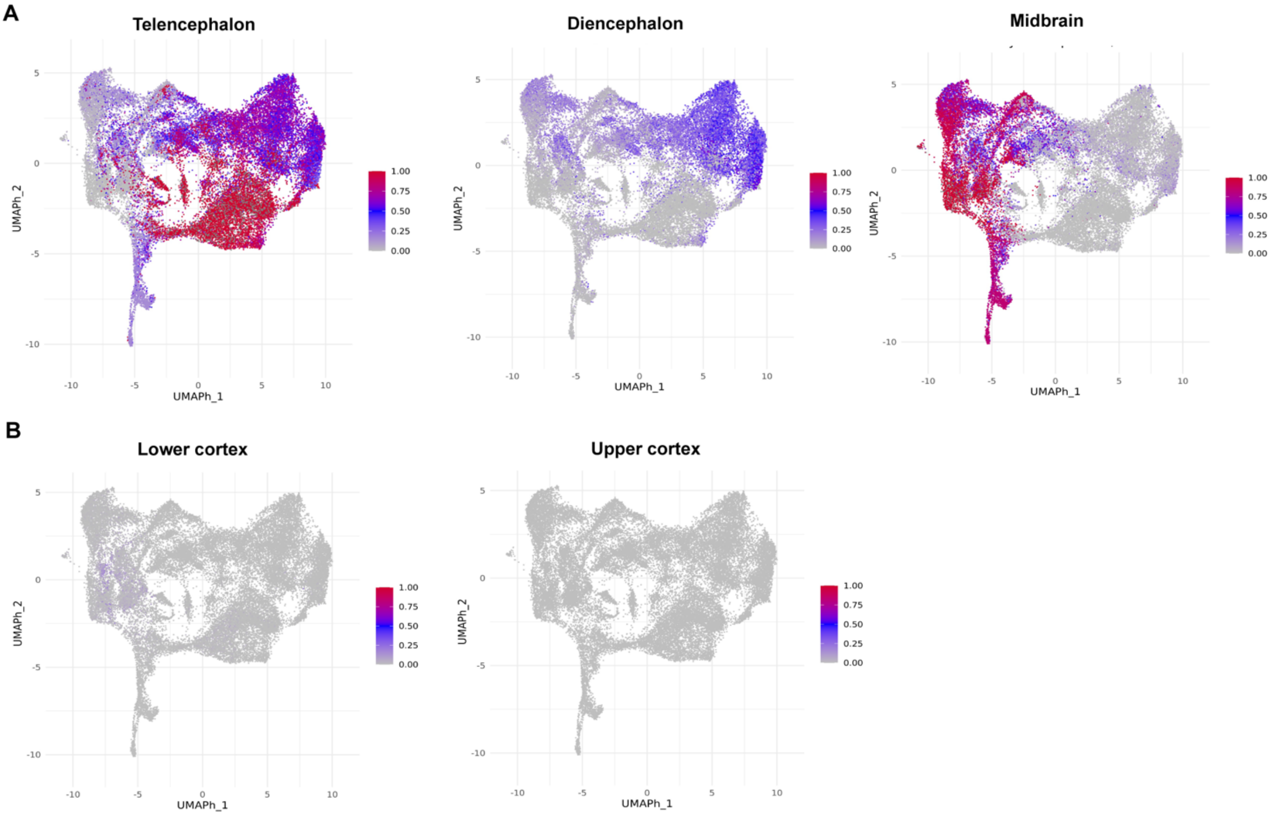


***Figure S5. Fetal brain atlas mapping of HD hOLS***

***(A)*** *UMAP representation of D150 HD hOLS mapped to the Braun fetal human brain atlas using cells from early developmental stages (5–7 post-conceptional weeks [pcw]). Predicted identity scores are represented by the color scale (grey, low; red, high).* ***(B)*** *UMAP representation of D150 HD hOLS mapped to the Braun fetal human brain atlas across the broader developmental range available for this analysis (5–14 pcw), including the stages shown in* ***(A)****, to assess mapping across developmental stages. Predicted identity scores are represented by the color scale (grey, low; red, high).*

### Supplementary Figure 6

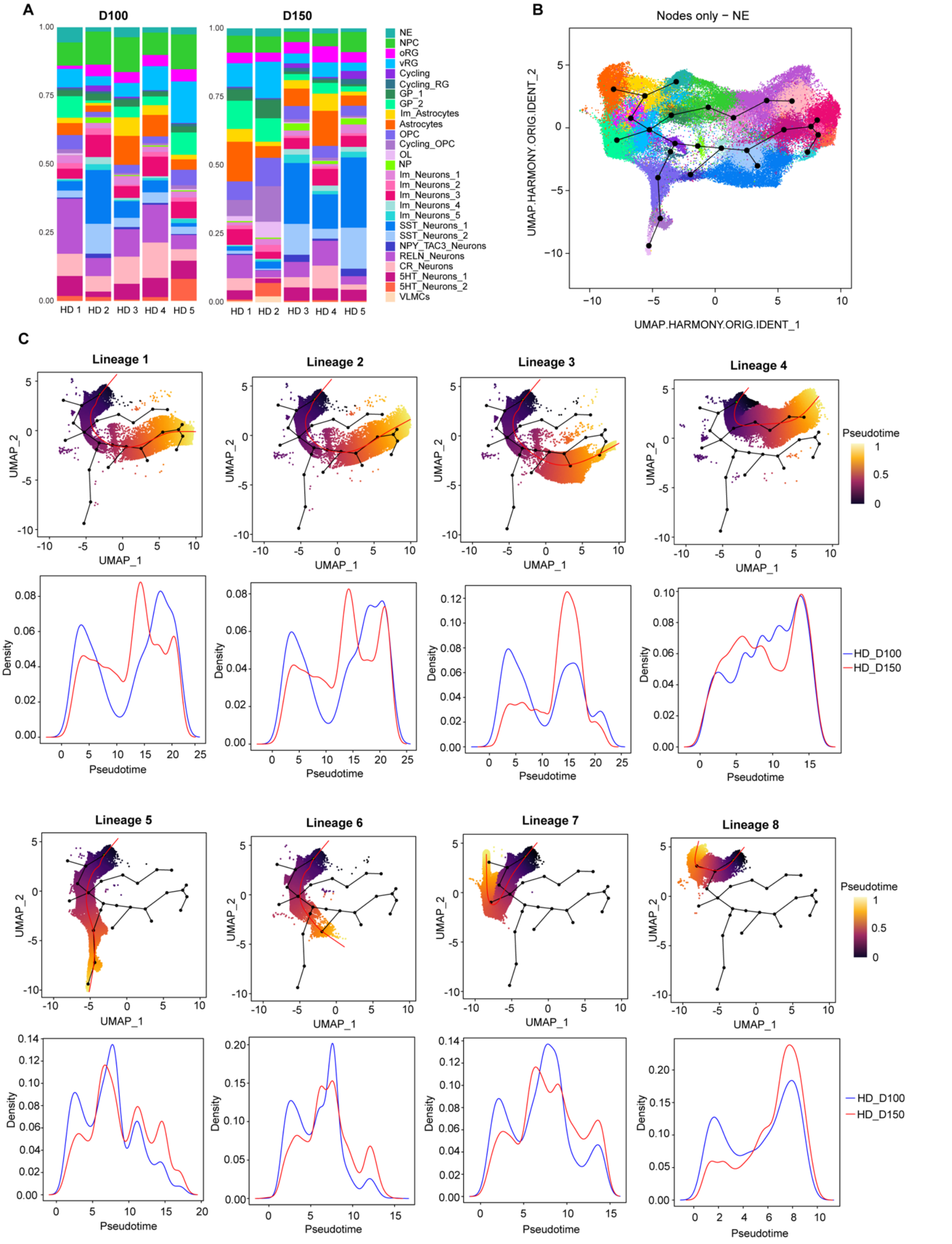


***Figure S6. Cellular composition and developmental trajectories of HD hOLS***

***(A)** Bar plots showing the proportions of cell clusters identified by scRNA-seq across HD samples (HD1–HD5) at D100 and D150. **(B)** UMAP representation of pseudotime analysis in HD hOLS. Nodes represent branch points along the inferred developmental trajectories, and lines indicate the reconstructed lineage paths. **(C)** Inferred lineage trajectories (Lineages 1–8)* *projected onto the UMAP. Corresponding density plots show the distribution of cells along each trajectory. Lineage 5 (oligodendrocytes) and Lineages 7 and 8 (astrocytes) are also shown in Fig. 3C. For cell-type abbreviations, see the legend to Fig. 2.*

**Supplementary Figure 7**

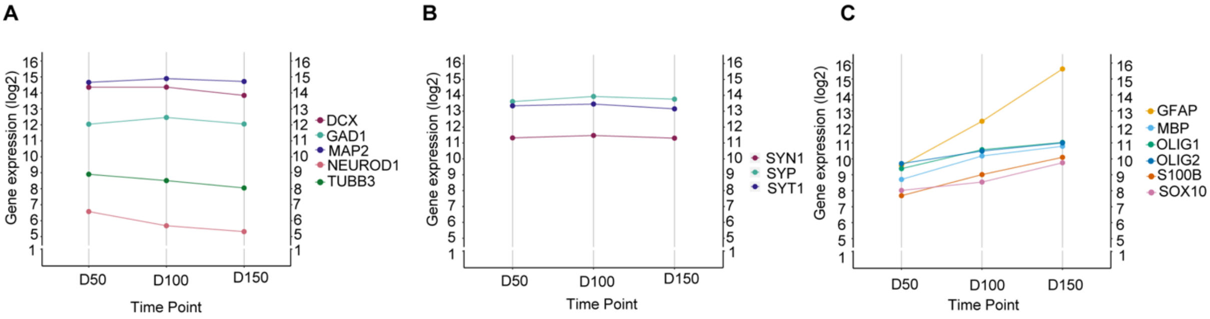


***Figure S7. Temporal expression dynamics of glial, neuronal, and synaptic markers in HD hOLS***

*Time-course expression of **(A)** neuronal genes (DCX, GAD1, MAP2, NEUROD1, TUBB3), **(B)** synaptic genes (SYN1, SYP, SYT1), and **(C)** glial genes (GFAP, MBP, OLIG1, OLIG2, S100B, SOX10), determined by bulk RNA-seq analysis of HD hOLS.*

### Supplementary Figure 8

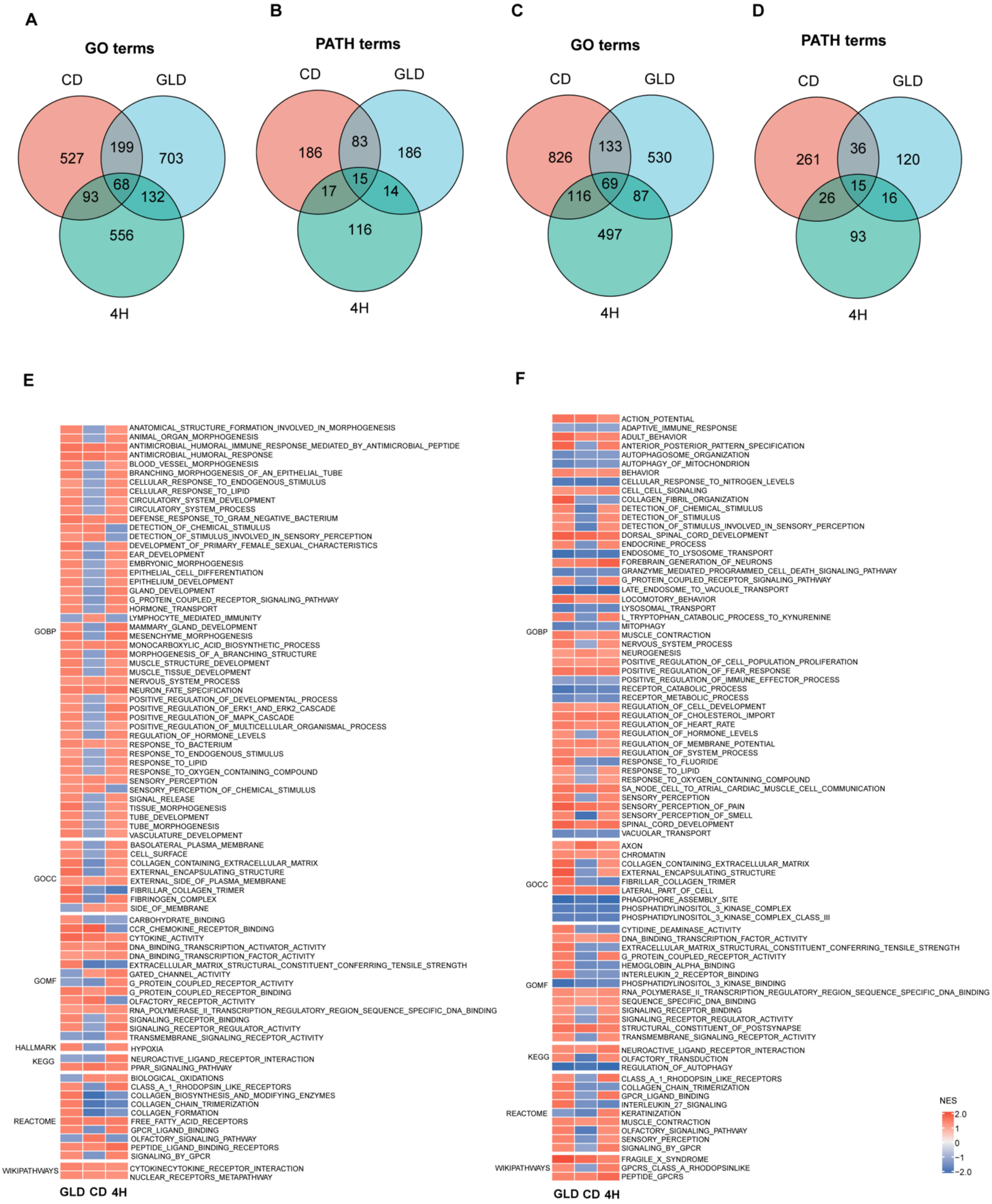


***Figure S8. Shared and disease-specific pathway alterations across leukodystrophy hOLS***

***(A–D)*** *Venn diagrams showing shared and disease-specific GSEA terms identified by bulk RNA-seq analysis of LKD hOLS at D100 and D150. GO terms are shown in* ***(A, C)*** *and pathway (PATH) terms in* ***(B, D)****.* ***(E, F)*** *Heatmaps showing normalized enrichment scores (NES) of GSEA terms commonly upregulated or downregulated across LKDs at D100* ***(E)*** *and D150* ***(F)****.*

### Supplementary Figure 9

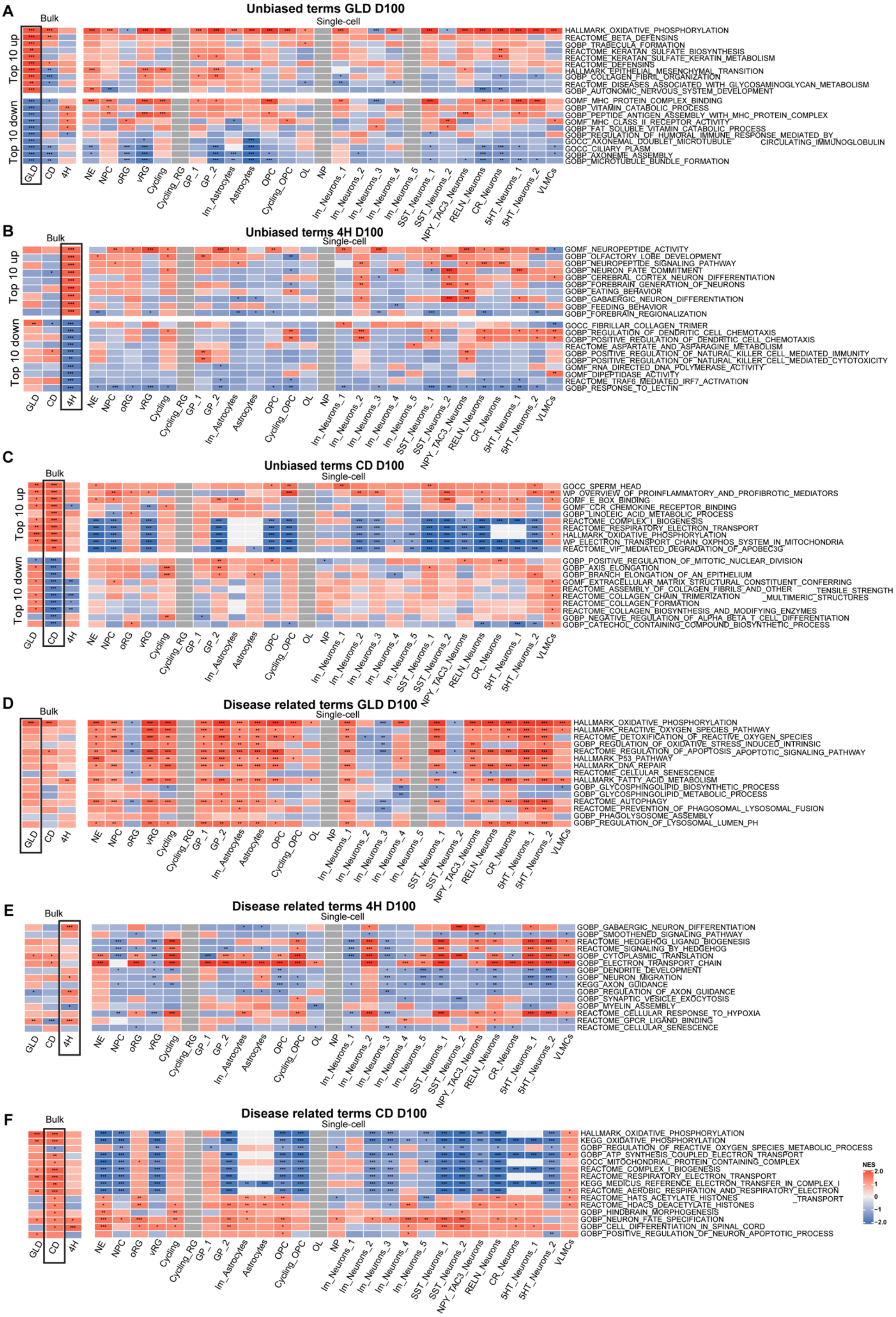

***Figure S9. Early pathway dysregulation in leukodystrophy hOLS at D100***

*Heatmaps showing normalized enrichment scores (NES) of GSEA terms identified by bulk RNA-seq and their enrichment across corresponding scRNA-seq cell populations in LKD versus HD hOLS at D100.* ***(A–C)*** *Top 10 upregulated and downregulated GSEA terms identified by bulk RNA-seq in GLD versus HD* ***(A)****, 4H versus HD* ***(B)****, and CD versus HD* ***(C)****; boxed columns indicate the respective bulk RNA-seq comparison used to select the terms.* ***(D–F)*** *Selected disease-relevant GSEA terms for GLD* ***(D)****, 4H* ***(E)****, and CD* ***(F)****, shown across bulk RNA-seq comparisons and the corresponding disease-versus-HD scRNA-seq cell populations. Statistical significance is indicated as follows: *p < 0.05, **p < 0.01, ***p < 0.001. For cell-type abbreviations, see the legend to Fig. 2.*

### Supplementary Figure 10

A

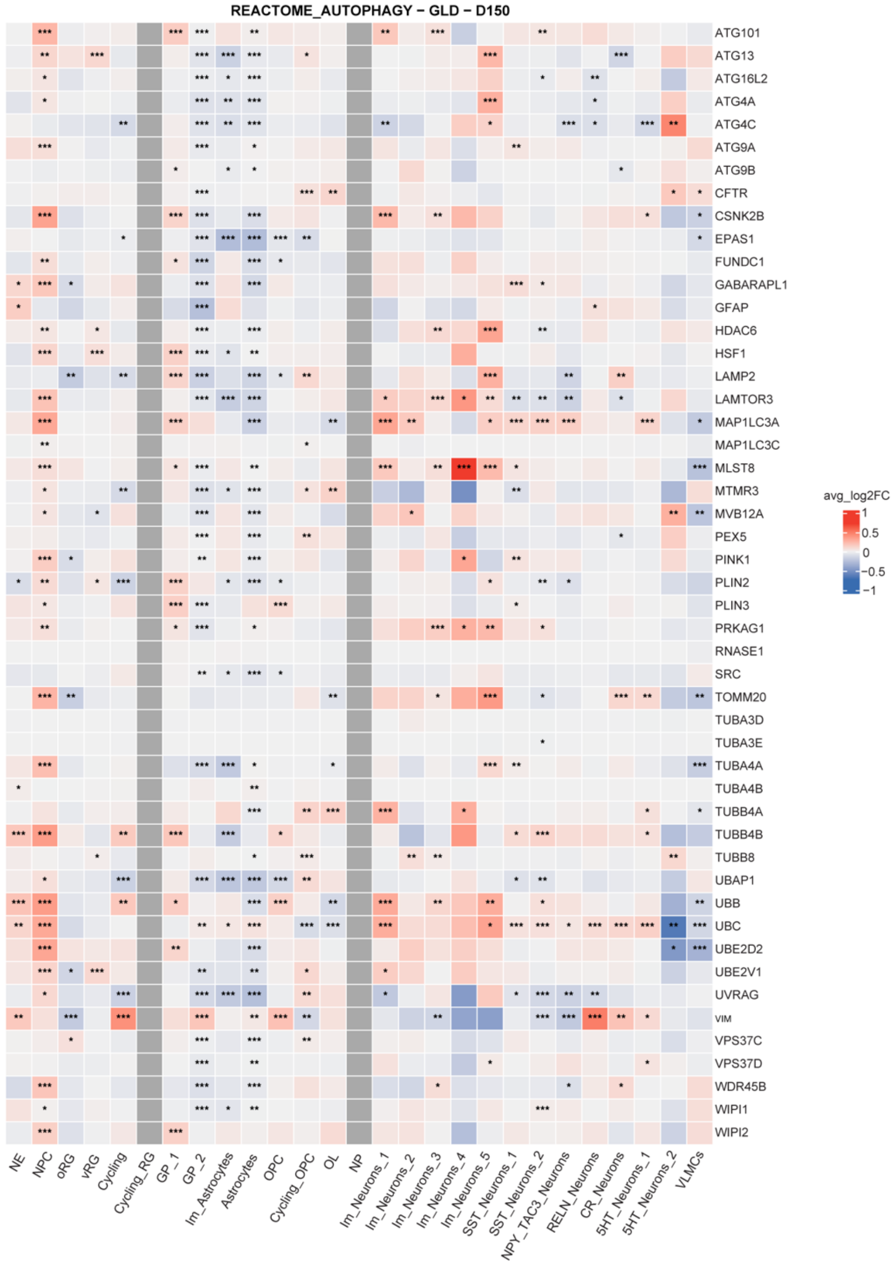


B

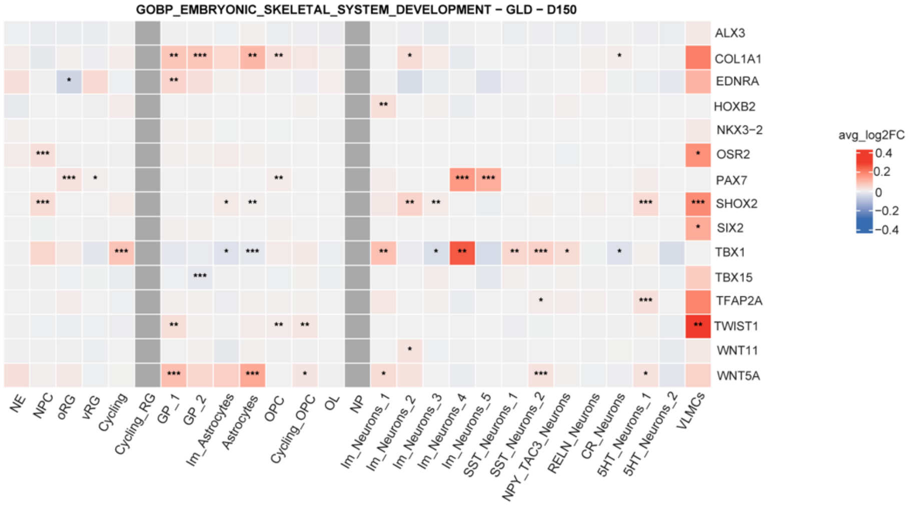

***Figure S10. Core enrichment genes of selected pathways in GLD hOLS at D150***

*Heatmaps showing core enrichment genes (rows) of selected GSEA terms across cell clusters (columns) identified by scRNA-seq analysis of GLD versus HD hOLS at D150. The color scale represents average gene expression (red, higher; blue, lower). Statistical significance is indicated as follows: *p < 0.05, **p < 0.01, ***p < 0.001.* ***(A)*** *REACTOME_AUTOPHAGY.* ***(B)*** *GOBP_EMBRYONIC_SKELETAL_SYSTEM_DEVELOPMENT. For cell-type abbreviations, see the legend to Fig. 2.*

### Supplementary Figure 11

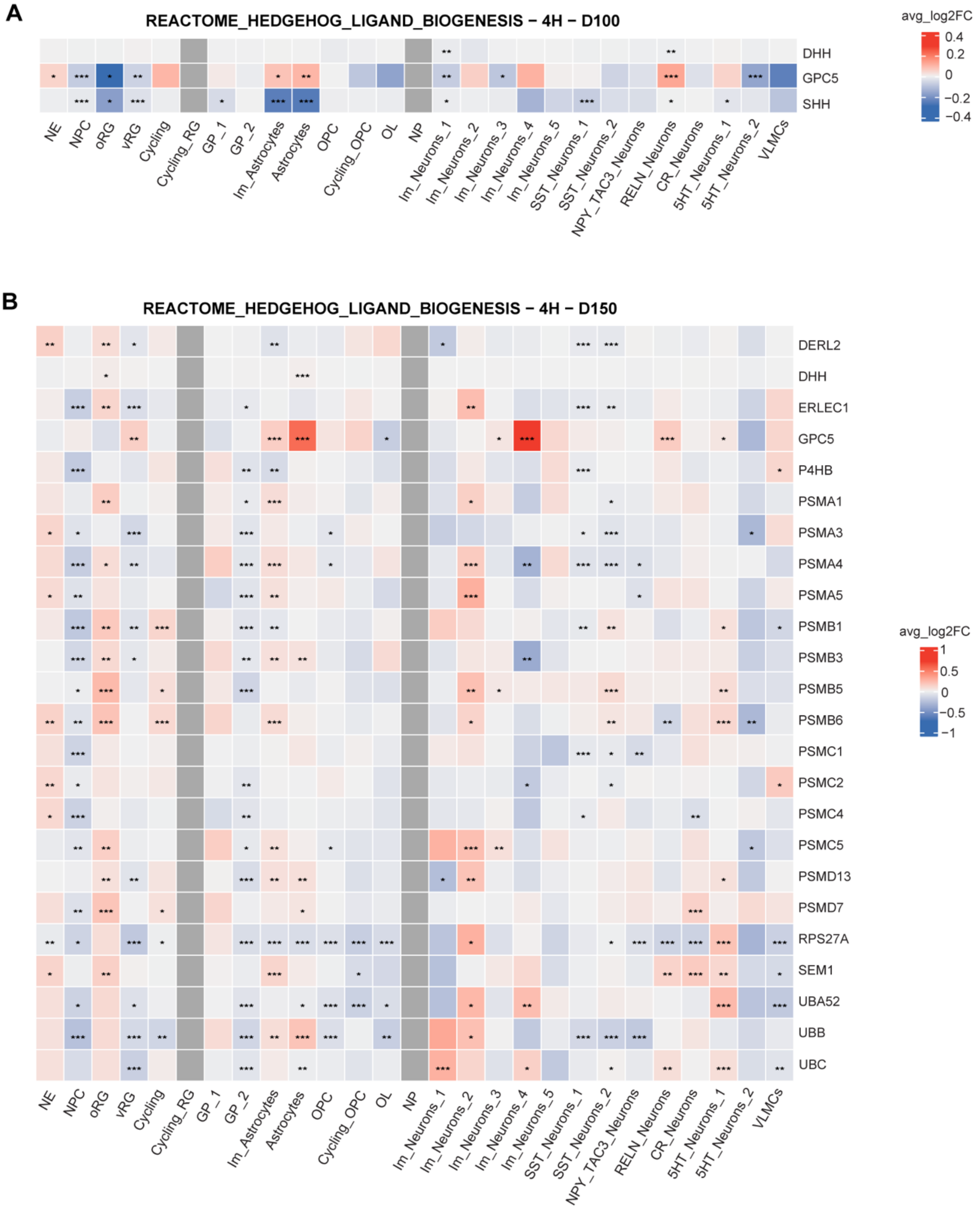


***Figure S11. Core enrichment genes of selected pathways in 4H hOLS at D100 and D150***
*Heatmaps showing core enrichment genes (rows) of selected GSEA terms across cell clusters (columns) identified by scRNA-seq analysis of 4H versus HD hOLS at **(A)** D100 and **(B)** D150. The color scale represents average gene expression (red, higher; blue, lower). Statistical significance is indicated as follows: *p < 0.05, **p < 0.01, ***p < 0.001. For cell-type abbreviations, see the legend to Fig. 2.*

### Supplementary Figure 12

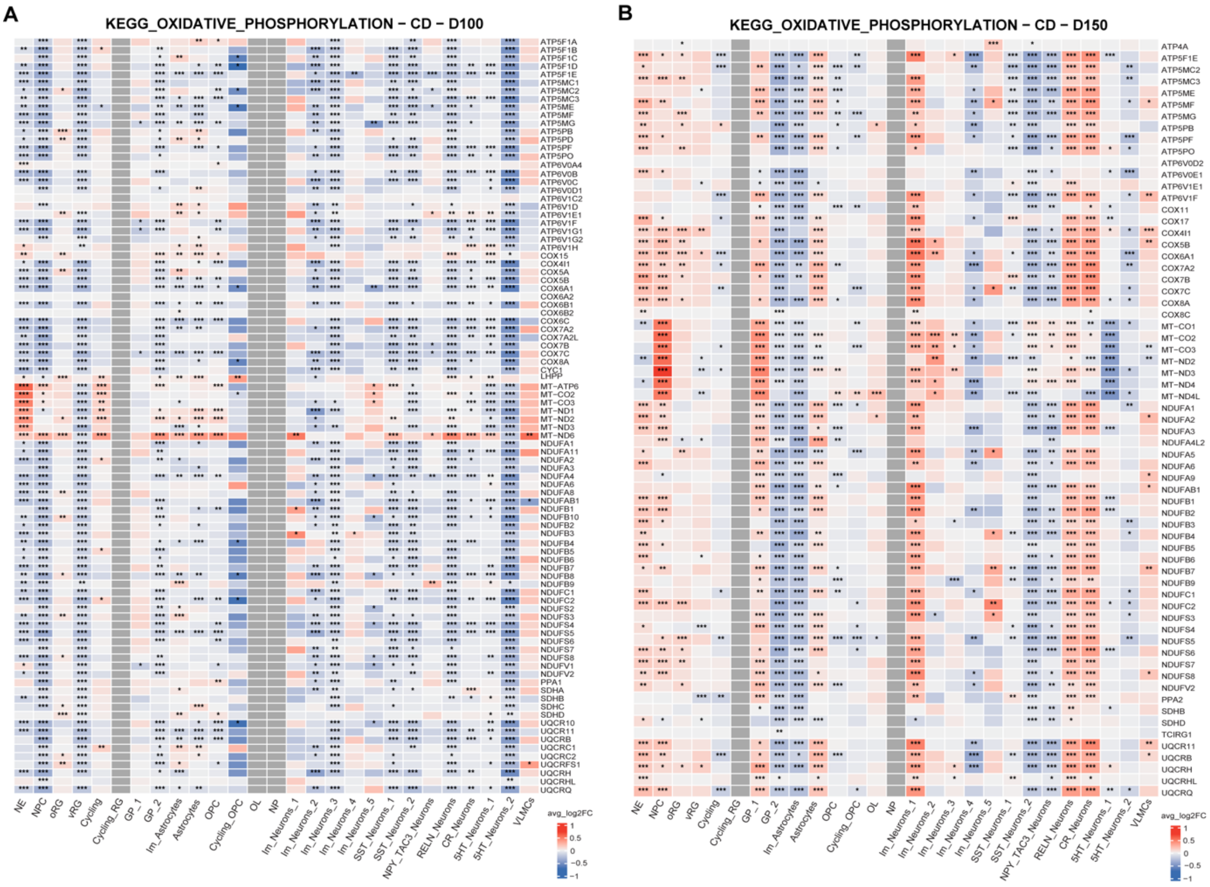


***Figure S12. Core enrichment genes of selected pathways in CD hOLS at D100 and D150***
*Heatmaps showing core enrichment genes (rows) of selected GSEA terms across cell clusters (columns) identified by scRNA-seq analysis of CD versus HD hOLS at **(A)** D100 and **(B)** D150. The color scale represents average gene expression (red, higher; blue, lower). Statistical significance is indicated as follows: *p < 0.05, **p < 0.01, ***p < 0.001. For cell-type abbreviations, see the legend to Fig. 2.*

### Supplementary Figure 13

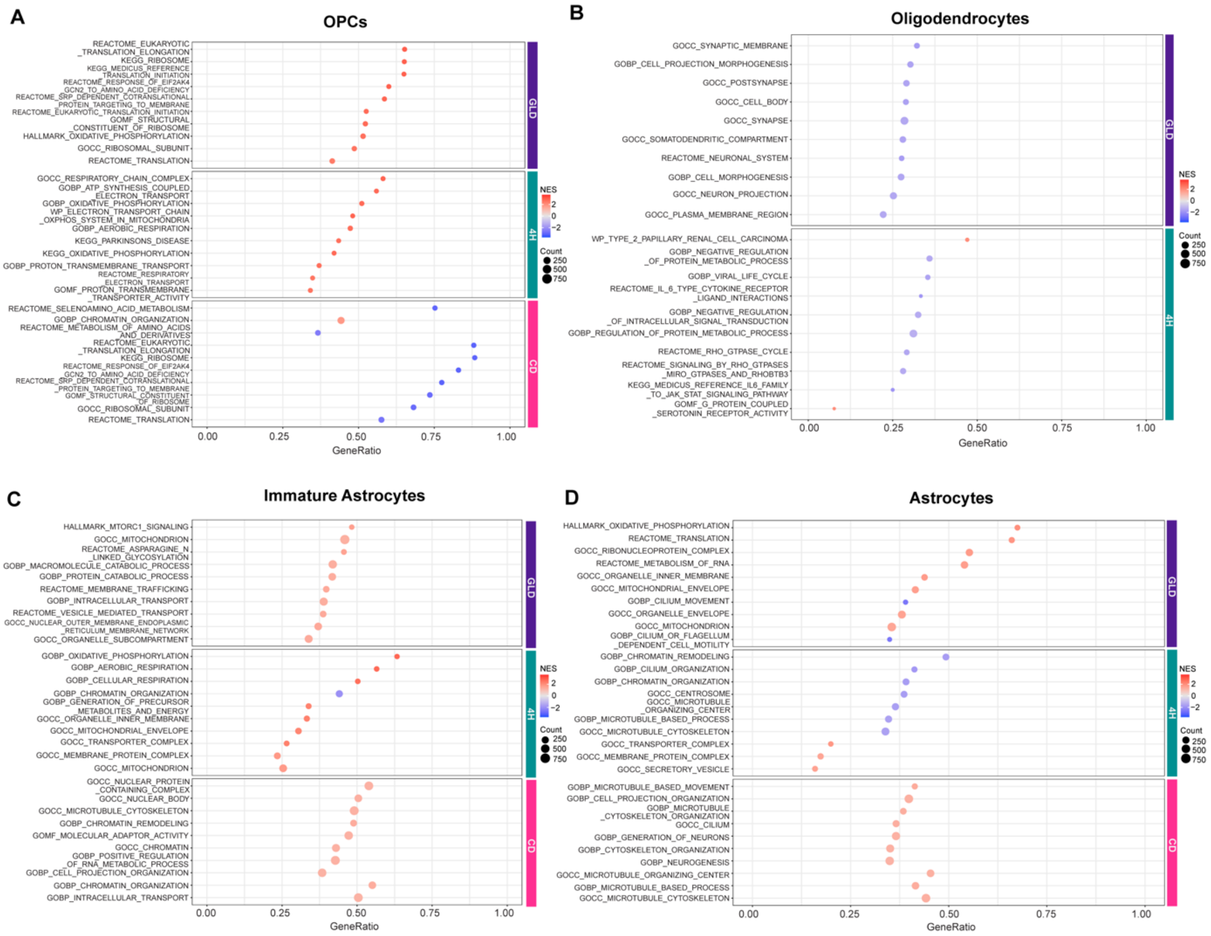


***Figure S13. Lineage-resolved pathway enrichment signatures at D100.***
*Bubble plots showing the top 10 enriched pathways identified by GSEA of scRNA-seq data from GLD, 4H, and CD hOLS compared with HD hOLS at D100 within **(A)** Oligodendrocytes Progenitors Cells (OPC), **(B)** Oligodendrocytes, **(C)** Immature Astrocytes, and **(D)** Astrocytes. Color indicates the normalized enrichment score (NES), and bubble size represents the number of genes contributing to each pathway.*

**Supplementary Table 1: Primary Antibodies**

| Target | Host | Dilution | Company | Number |
|---|---|---|---|---|
| GABA | Rabbit | 1:300 | Sigma | A2052 |
| GFAP | Mouse | 1:1000 | Millipore | MAB3402 |
| MBP | Rat | 1:300 | Chemicon | MAB386 |
| NF | Mouse | 1:300 | Sigma | N0142 |
| Olig2 | Rabbit | 1:200 | Abcam | ab136253 |
| TUBB3 | Mouse | 1:1000 | Biolegend | 801202 |
| MAP2 | Mouse | 1:700 | Millipore | MAB3418 |
| VIMENTIN | Mouse | 1:200 | Merck | V5255 |
| FABP7 | Rabbit | 1:300 | Millipore | ABN14 |
| TPH2 | Rabbit | 1:100 | Proteintech | 29283-1-AP |
| NPY | Rabbit | 1:100 | Cell Signaling | 11976 |

**Supplementary Table 2: Secondary Antibodies**

| Target | Host | Dilution |
|---|---|---|
| Alexa 488 | Goat anti-mouse IgG (Mol.Probes, A11001) | 1:1000 |
| | Goat anti-rabbit IgG (Mol.Probes, A11008) | 1:1000 |
| | Goat anti-rat IgG (Mol.Probes, A11006) | 1:1000 |
| Alexa 546 | Goat anti-mouse IgG (Mol.Probes, A11003) | 1:2000 |
| | Goat anti-rabbit IgG (Mol.Probes, A11010) | 1:2000 |
| Alexa 647 | Goat anti-rabbit IgG (Mol.Probes, A21246)<br>Goat anti-mouse IgG (Mol.probes, A21235) | 1:500 |
